# Apidae BeeHavioural response to shape-neutral visual stimuli in natural setup

**DOI:** 10.64898/2026.02.06.702584

**Authors:** Josi Ruschinczyk, Selina Braungart, Paul Hertel, Carolina Benkewitz, Pedram Jalali

## Abstract

Floral displays attract pollinators through a finely tuned interplay of colour, pattern, shape, and scent. Yet, the question remains: how do bees respond when these traits are stripped to their simplest form, with only visual cues at play? In this field study, we examined the foraging behaviour of Apis mellifera on artificial flowers differing solely in background colour (white or yellow) and UV patterning, while shape and scent were held constant. Across three summer days, standardized stimuli were placed within a natural meadow, and bee–flower interactions were recorded and analyzed by Bayesian hierarchical models. The results reveal a clear preference for yellow over white backgrounds and prolonged visitation in the presence of ring-shaped UV patterns, whereas full UV coverage acted as a deterrent. These effects, though moderate, were consistently modulated by abiotic covariates, particularly radiation, temperature, and time of day. Negligible inter-individual variation and a substantial share of residual variance further underline the context-dependent complexity of foraging. In sum, our findings demonstrate that visual floral traits, while influential, are interpreted through the dual lens of environmental contingency and the bees’ inherent cognitive machinery.

## Introduction

The interplay of sensory abilities in bees (Apidae) forms the foundation of their role as key pollinators in ecosystems. Their interactions with flowers represent an intricate, yet vital ecological relationship (Kearns et al., 1998) which is controlled by an assortment of sensory organs and has a significant impact on their foraging efficiency. Among their less commonly recognized but noteworthy modalities are bees” electric-field sensitivity (Hsu et al., 2007), their inherent time perception which directly regulates their foraging behaviour (Moore et al., 1989). Bees also have developed tactile discernment that allows them to distinguish between different surface textures and floral parts (Scheiner et al., 2005).

Nevertheless, olfactory and visual cues remain the most prominent sensory path among members of Apidae. Their olfactory and taste receptors enable them to assess nectar quality (Liao et al., 2023) and its relative sugar content (Değirmenci et al., 2023; Haupt, 2004). Yet, it is the visual perception that helps the bees to perceive their surroundings, including movements, shapes and colour. This dual reliance on smell and vision illustrates the evolutionary advantage of integrated multisensory channels, providing the bees with necessary means of rapid and accurate decision-making (Kulahci et al., 2008).

The visual cue of *Apis mellifera* is divided between different eye structures; two large compound eyes (oculi) located dorsally on their head, and three simple eyes (ocelli) that occupy the ocular bridge in a triangular conformation. The ocelli are tasked with light intensity and shadow movement detection, while the colour vision is governed by oculi. Bees are trichromatic and rely on three types of photoreceptors (opsins) that are unevenly distributed among the rhabdoms of somewhat 5,000 to 6,000 omatidia which form the compound eyes; yet, unlike humans, the Apidae opsins are responsive to shorter wavelength spectrum with maximum sensitivity of: 540 nm in the green range, 430 nm in blue and 340 nm in the UV-A (R. Menzel & Blakers, 1976; Peitsch et al., 1992). The actual colour is believed to be perceived by opponent-process theory, which can be represented in a hexagon. This hexagon depicts the maximum stimulation of each photoreceptor and their opposites in order to optimize a tangible color system through the relative excitation of the opposites (Briscoe & Chittka, 2001). The evolutionary roots of colour-vision in bees can be traced back to the Devonian ancestors of winged insects and is the evidence of insect-flowering plants co-evolution (Chittka, 2017).

Bee foraging behaviour varies with flower shape and colour. Naive foragers rely heavily on morphological traits such as shape and symmetry, while the foraging of experienced ones is heavily weighted by olfactory signals (Orbán & Plowright, 2014). Flowers with high spectral purity (saturation) in the bees” visual range in addition to radial or concentric symmetry are generally preferred by Apidae. Importantly, bee colours do not map directly onto human colour perception (R. Menzel & Blakers, 1976; Peitsch et al., 1992). In particular, UV patterns/patches, which are often complemented by yellow (green + UV) or violet (blue + UV) floral ground colouring, enhance the visual contrast and promote the attractiveness of the flowers (Guldberg & Atsatt, 1975; Ibarra et al., 2022). These contrasts facilitate the locating of individual flowers and act as nectar guides, making access to the rewards more efficient (Chittka & Raine, 2006). Furthermore, flower colors are evolutionarily adapted to the spectral sensitivity of pollinators, as evident via pollinator-driven selection processes, revealing that flowers in different habitats develop convergently similar color distributions (Chittka & Menzel, 1992; Dyer et al., 2012). Additionally, development of similar UV-patterns can be clearly demonstrated by UV photography (Koski & Ashman, 2014). Indeed, comparative analyses show that plant species with distinct UV patterns tend to achieve higher pollination rates (Koski & Ashman, 2015).

As flowers are complex and dynamic organs and their nature makes it rather impossible to discern the reaction to visual cues from other modalities, many ethological experiments turned to artificial flowers or dummies. Artificial flowers allow for specific variable traits and make it possible to individually address their effect on cognitive chain activation behind the flower choice (Chittka & Wells, 2004). One of the main outcomes of this approach is the capability of bees at learning complex patterns and linking color contrasts with a reward (Giurfa et al., 1996; Lehrer et al., 1995). Furthermore, as artificial flowers make it possible to evaluate (re)cognitive performance and similar colour generalization effect, they are the best tools in assessments of learning process and memory handling behind bees” food selection, (Dyer & Chittka, 2004; Giurfa & Vorobyev, 1997). The common outcome of these controlled approaches is that distinct color patterns lead to increased attractiveness. However, most of these studies were conducted under enclosure in laboratory environments, omitting subtle yet important parameters, such as background vegetation”s effect, natural light component and daily/seasonal intensity variation, and the competition between scent and shape signals; therefore, their result cannot be simply generalized to *in vivo* state. This methodological limitation calls for reflection on the transferability of the results to ecological contexts (Chapman et al., 2023; Ibarra et al., 2022).

While numerous studies showed that color contrast leads to increased attractiveness, the response of pollinators to artificial flowers without variables such as accompanying colour, scent and shape, remains ambiguous; as in natural systems, these traits almost always overlap and reinforce or attenuate each other (Kaczorowski et al., 2005; Raguso, 2004). A prominent instance is the influence of olfactory cues on the visual patterns learning rate (Leonard et al., 2010), or interference of certain flowers in color contrast effectiveness (Ibarra et al., 2015). The present study is an endeavor to assess the cognitive responses of Apidae to artificial flowers with controlled visual traits, yet in a natural setup—aiming to reach more conclusive results that retain their validity amid biological and ecological noise.

## Materials and Methods

The experiment was carried out in three chronological replicates (19–21 August 2025) in an open meadow, with fully randomized experimental plot selection and semi-randomized stimulus distribution to mimic the stochastic arrangement of flowers in nature. Four UV pattern allocations on two background colours were set as experimental variables, with five local replicates per treatment per day, while all other parameters were held neutrally constant.

The meadow was located at the test site of the Fraunhofer-Institute (51.0266563 N, 13.7388663 E; Räcknitz, Dresden-Plauen, Germany). It contained a mixture of native flowering plants and was last mowed on 1 July 2025, with an average vegetation height of ∼60 cm at the time of the experiment. During the experiment, two commercial honeybee colonies were located on the meadow, approximately 55,66 m away from the experimental plot (linear distance to the plot center).

The artificial flowers were designed following similar principles to Russell & Papaj (2016) and Chapman et al. (2023) (Chapman et al., 2023; Russell & Papaj, 2016). To assemble, decapped 1.5 ml centrifuge tubes, serving as nectar reservoirs, were inserted through the center of paper rings (ø 3.0 cm, rim width 2 cm). Rings for the white background were cut from Canson Bristol paper (250 g/m^2^), and those for the yellow background from URSUS construction paper (130 g/m^2^, Zitronengelb), and labeled on the reverse side. Each reservoir was fitted with a cotton wick, consisting of a narrow tail and a bulky dome-shaped stopper that protruded above the tube to allow nectar transfer.

The UV patterns were applied daily prior to the experiment by spraying the designated sections with a commercially available transparent sunscreen spray (Eucerin® Sun Protection, 50+ SPF, Oil Control; active substances: Diethylamino Hydroxybenzoyl Hexyl Benzoate, Ethylhexyl Salicylate). For stimulus A (ring), the entire paper ring was treated; for stimulus B (core), only the dome-shaped cotton wick in the center was sprayed; for stimulus C (full), both the paper ring and the central wick were treated, while stimulus D (null) remained untreated.

The pattern application was validated using UV imaging with a Sony Alpha NEX-5T mirrorless camera (Quartz sensor window) equipped with a Sigma 30 mm f/2.8 DN Art lens, a No. 2 Kenko close-up filter, and a Baader Venus U-filter.

The assembled artificial flowers were mounted on approximately 60 cm long bamboo sticks (ø ≥ 0.7 cm) using rubber bands. Reservoirs were filled to the 1.0 ml mark with freshly thawed aliquots of artificial nectar (prepared on 18 August and stored at −20 °C), consisting of 30% (w/v) sucrose, 0.6% (v/v) linalool, and 0.1% (v/v) phenylacetaldehyde.

Plot randomization was achieved by throwing an ecological sampling quadrat (50 × 50 cm, subdivided into 100 cells of 5 × 5 cm). Vegetation within the selected quadrat was flattened to reduce competition. The 40 stimuli were then distributed under the following constraints: (i) no more than 20 cells contained a single stimulus, (ii) cells could contain two stimuli of either the same or different colours, (iii) up to three stimuli were allowed per cell if they were either all of the same colour or a combination of two of one colour plus one of another, and (iv) within any 3 × 3 block of cells, no more than six stimuli of the same colour were permitted.

**Figure 1.**
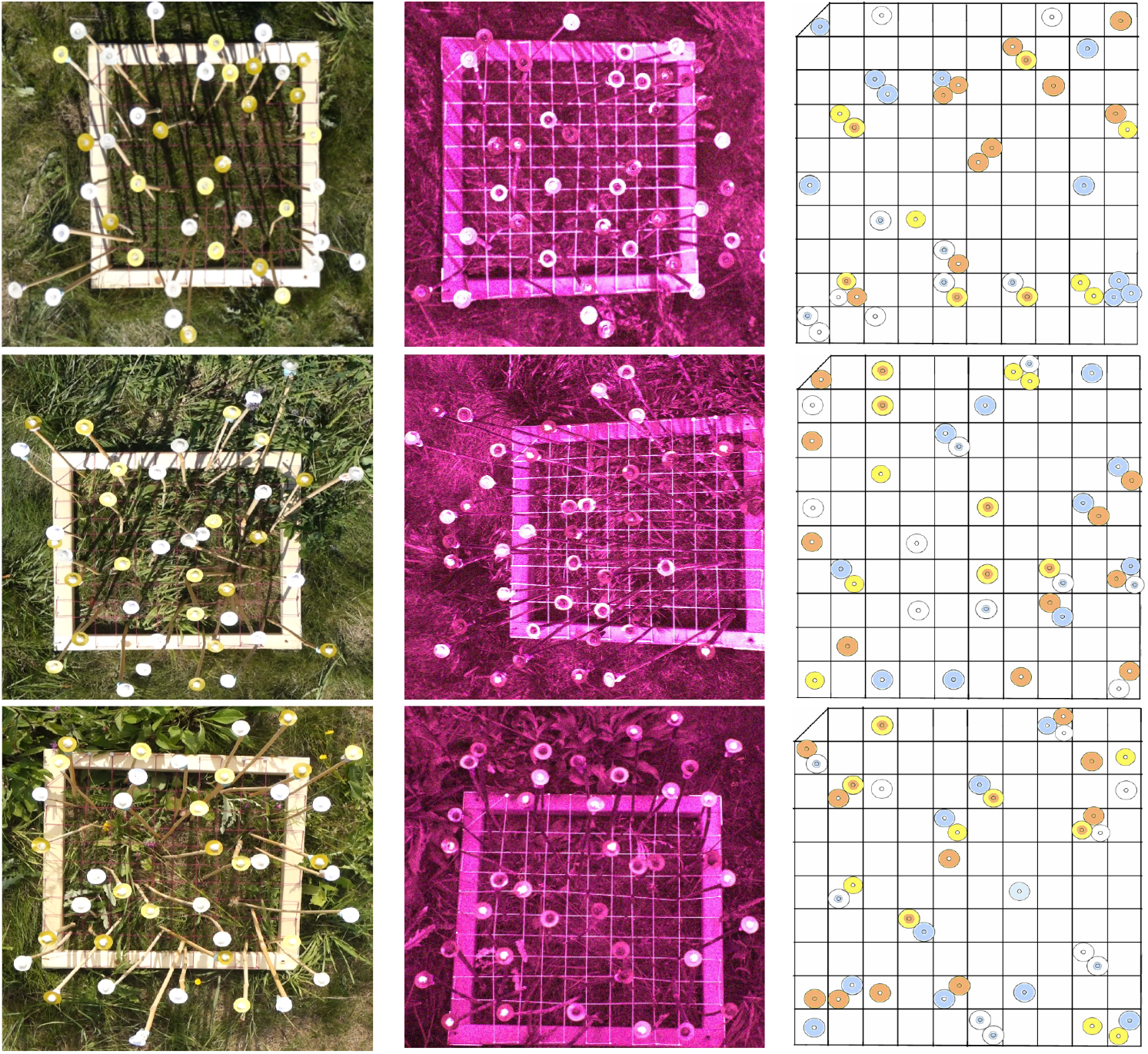
Map of artificial distribution within each experimental day. (A) Stimuli distribution on Tuesday (August 19, 2025). (B) Stimuli distribution on Wednesday (August 20, 2025). (C) Stimuli distribution on Thursday (August 21, 2025).

A Sony Cyber-shot DSC-RX100M3 camera was mounted horizontally above the experimental plot on a Wallimex WT-501 boom stand, and continuous 3-hour recordings of pollinator interactions with artificial flowers were made between 10:00 and 14:00.

All video recordings from the three experimental days were surveyed independently by four observers. Observers used a time-stamped, event-based evaluation sheet to document pollinator activity. For each visit, the observer assigned a Visitor identity (VisitorID) to the individual pollinator and recorded the event type (landing or take-off) on a given artificial flower (FlowerID). To ensure consistency, each VisitorID was retained for as long as the same individual remained within the recording frame and reassigned only when a pollinator left the frame and re-entered. The family and genus of the visitor were noted in the Category column. Timestamps were extracted by Shotcut software (Meltytech, LLC, 2025). Each observer completed the full survey of all three experimental days individually, ensuring complete replication of annotations across observers.

Environmental parameters for Dresden were obtained daily from the Deutscher Wetterdienst (German Weather Service) using the official weather station in Dresden-Klotzsche.

**Figure 2.**
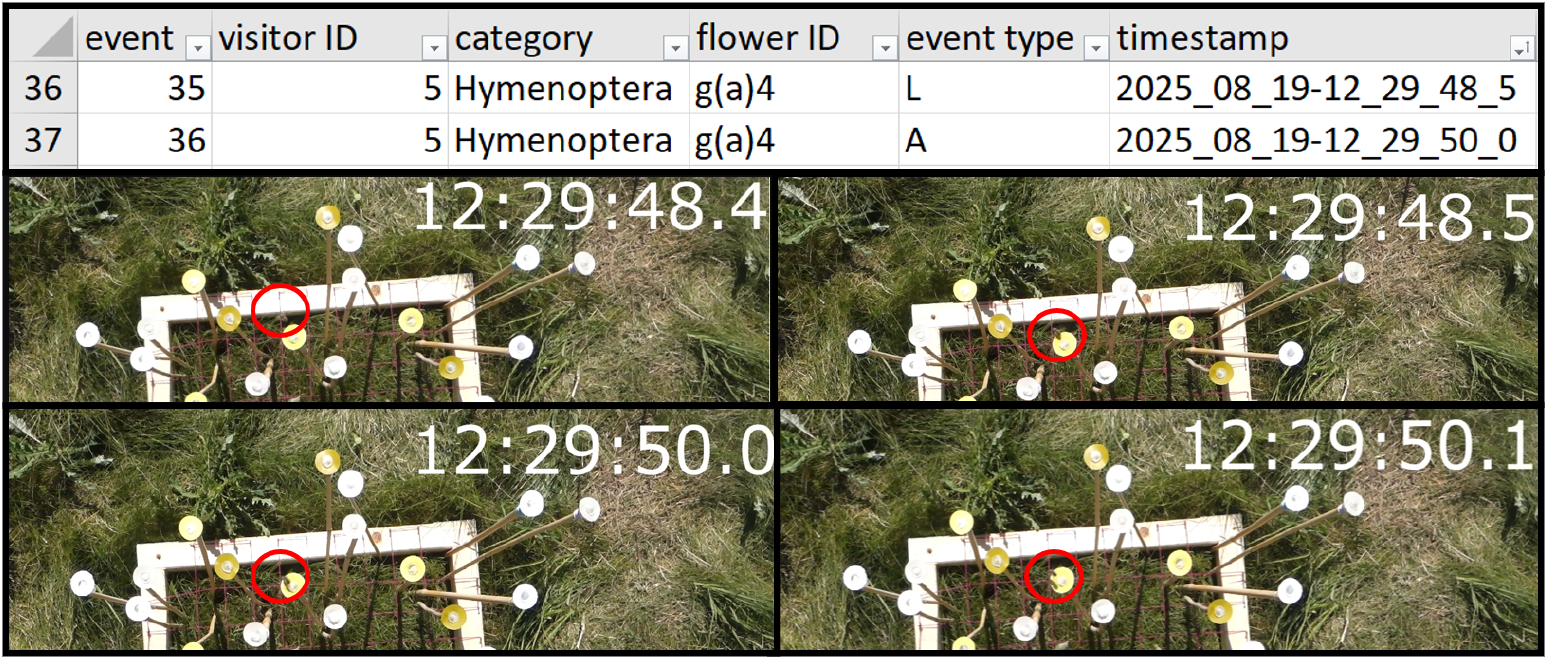
Shown is an excerpt from an observer”s formatted table with two events for a visitor ID. Below are four screenshots from the specified day, showing the visitor on approach, landing, before, and during takeoff. The visitor is marked with a red circle.

Prior to statistical modeling, the dataset was filtered to exclude all non-Apidae visitors and incomplete events (i.e., landings without a corresponding take-off, or vice versa). Weather variables, recorded at 10-minute intervals, were linearly interpolated to preserve their continuous structure. Each visit event was then linked to the corresponding standardized (z) value of the weather covariates: mean wind speed, air temperature at 2 m, relative humidity, global solar radiation, and long-wave downwelling radiation. A minimum dwell time of 3 s was defined as the threshold for foraging, with foraging itself interpreted as the initiation of the visual cognitive response chain.

To evaluate the effects of background colour and UV pattern allocation on both the probability of foraging and the duration of dwelling, Bayesian hierarchical regression models were applied. Posterior inference was obtained via Hamiltonian Monte Carlo sampling using the No-U-Turn Sampler (NUTS). Each model was run with four parallel chains, including 1,000 warm-up (tuning) iterations and 1,000 posterior draws per chain, resulting in 4,000 effective samples.

In both models, background colour, UV pattern allocation and their interaction, a cubic B-spline of experimental time (df = 4 over the 0–3 h session), the z-standardized experimental day (chronological replicate/learning effect), and z-standardized weather covariates (air temperature at 2 m, wind speed, relative humidity, global radiation, and long-wave downwelling radiation) were included as fixed effects. VisitorID, FlowerID, and observer bias were included as hierarchical random effects. Priors for fixed-effect coefficients were specified as weakly informative normal distributions, while priors for group-level (random-effect) standard deviations followed half-normal distributions. Both models used white background × null UV pattern as the baseline intercept to enable pairwise contrasts.

Foraging probability was estimated using a Bayesian logistic regression with a Bernoulli likelihood and logit link. Expected dwelling time was analyzed with a Bayesian accelerated failure-time (AFT) model assuming a log-normal likelihood for dwell time (in seconds).

Additionally, to characterize how abiotic light conditions modulate the effectiveness of visual floral cues, an extended foraging model was fitted that included treatment-specific quadratic effects of global radiation. This model retained the original fixed and random-effect structure but added linear and quadratic global radiation terms interacting with background colour and UV-pattern allocation. Posterior predictions from this model were then evaluated across the observed range of global radiation to estimate condition-specific radiation levels associated with maximum foraging probability.

Models reliability was assessed through posterior predictive checks, comparing synthetic data with the observed outcomes.

All analyses were performed in Python (v3.12)(Oliphant, 2007) within the Anaconda distribution, using Jupyter Notebook (Thomas et al., 2016) as the interactive environment. Bayesian models were implemented in PyMC (v5) (Patil et al., 2010) with sampling carried out by the No-U-Turn Sampler (NUTS). Model summaries and diagnostics were obtained with ArviZ (v0.16), and design matrices were constructed using patsy. Data manipulation and visualization relied on pandas, NumPy (Harris et al., 2020), and matplotlib (Hunter, 2007). The full computational workflow was managed in a reproducible pipeline within Jupyter Notebook, ensuring transparency and replicability.

## Results

Across all recordings, a mean total of 652.2±52.7 v events were annotated, which resulted in average of 326±19.1 paired events before and 323±18.7 events after filtering. The total of 3 excluded events where the visits from non-Apidae.

Both the foraging (Bernoulli–logit) and dwelling-time (log-normal AFT) models demonstrated excellent convergence and fit. Across all parameters, the Gelman–Rubin statistic 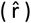 was equal to 1.00, effective sample sizes (ESS) exceeded 1,000 for both bulk and tail estimates, and no divergent transitions were reported. Posterior predictive checks indicated strong agreement between observed and model-simulated data, supporting the adequacy of model specification.

**Figure 3.**
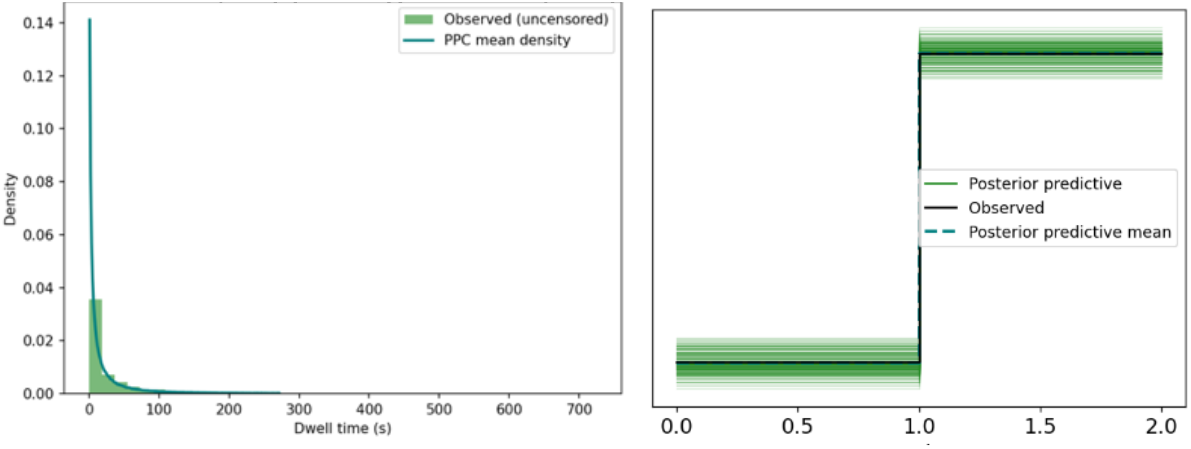
PPC of dwelling.time (le1) and foraging (right) model, revealing a very good agreement between posterior simulated and observed data.

The Bernoulli/logit foraging model showed no effects of background colour or UV allocation at the 95% HDI. At the 90% HDI, however, flowers with a yellow background were 27% more likely to be foraged than those with a white background, and flowers with a full UV pattern were 18.5% less likely to be foraged than those without UV.

**Table 1.**
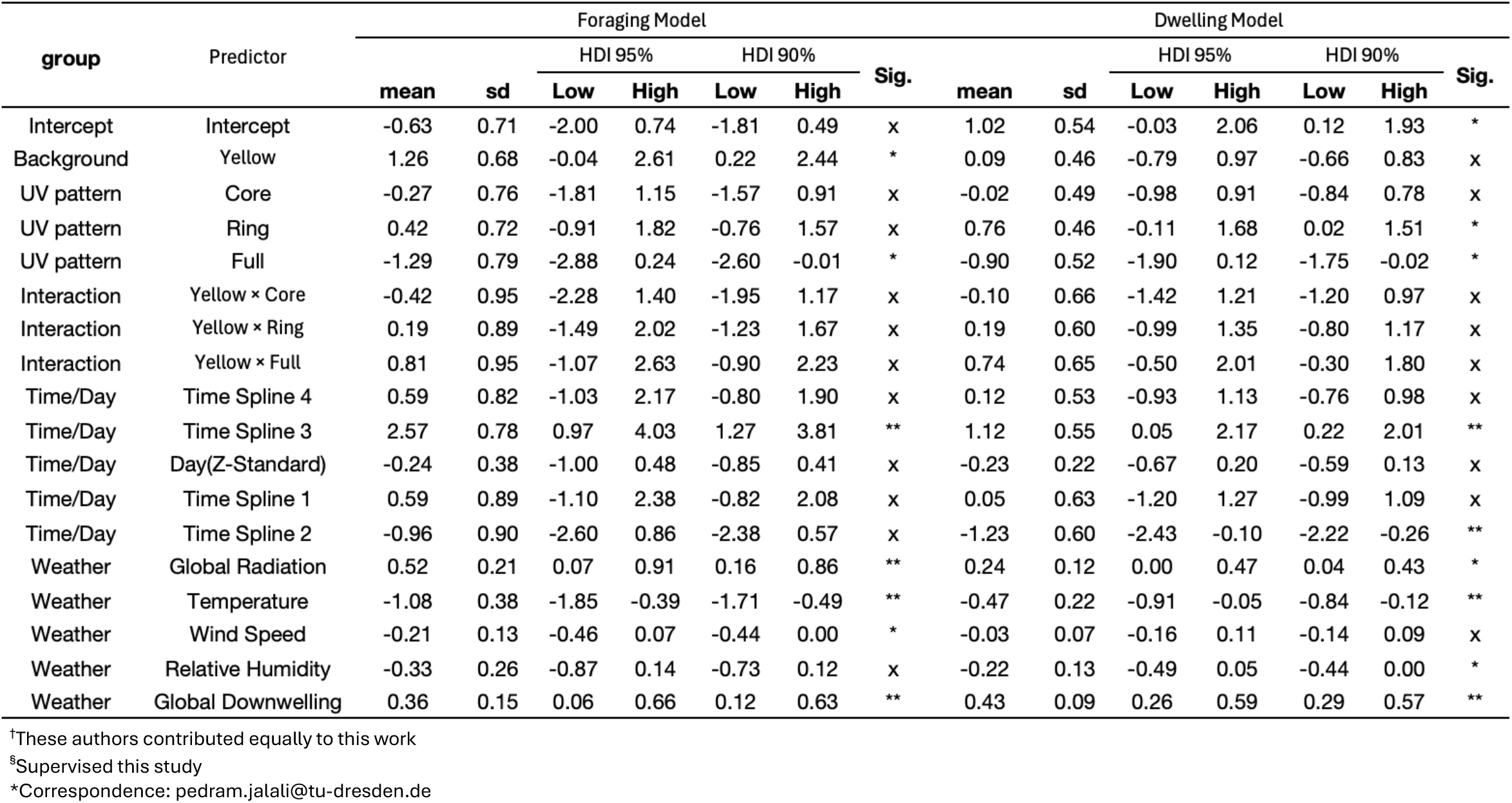
Log odds of foraging and log times of dwelling within fixed and random effects. ** indicate significance at 95% HDI, * at 90% HDI.

In the AFT/log-normal model, the intercept was significant at the 95% HDI, indicating that even under baseline conditions (white background, no UV pattern, average weather), dwell times were systematically longer than expected under a neutral model. Specifically, baseline foraging events were estimated to last up to ∼2.8 times longer, highlighting an inherent structure in dwelling behaviour independent of treatment effects. Neither background colour nor UV pattern allocation showed significant effects at the 95% HDI. However, at the more permissive 90% HDI, dwell times were likely ∼2.1 times longer on flowers with ring-shaped UV allocation, while flowers with full UV coverage were associated with ∼0.6 times shorter dwell times compared to no UV pattern at all.

The third time-spline (representing the smooth trend over the midday quartile) was significant at the 95% HDI in both the foraging and dwelling models, increasing the probability of foraging by ∼87% and extending dwell times by approximately 3.1-fold. The second time-spline (spanning late morning into midday) reached significance only in the dwelling model (95% HDI), where it shortened dwell times to ∼0.7-fold of baseline. The remaining spline terms crossed zero in both the 90% and 95% HDIs and were therefore considered non-significant. Notably, the monotone fixed effect of experimental day (the chronological replicate/learning effect) was also non-significant in both models.

**Figure 4.**
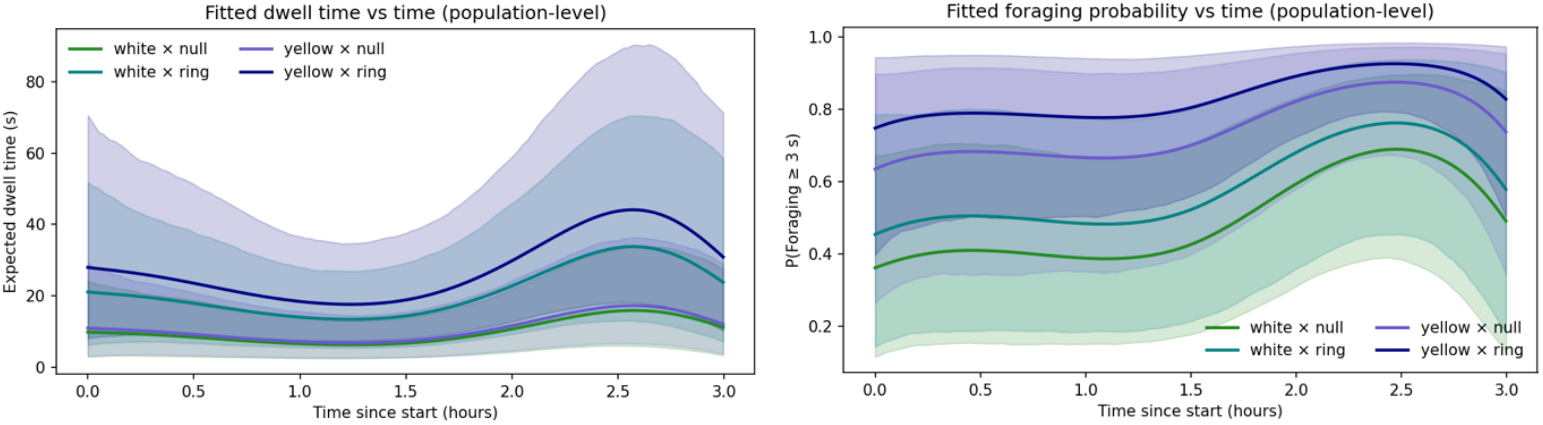
Distribution of foraging probability (right) and dwelling time (le1) among the four overlapping time-splines.

**Figure 5.**
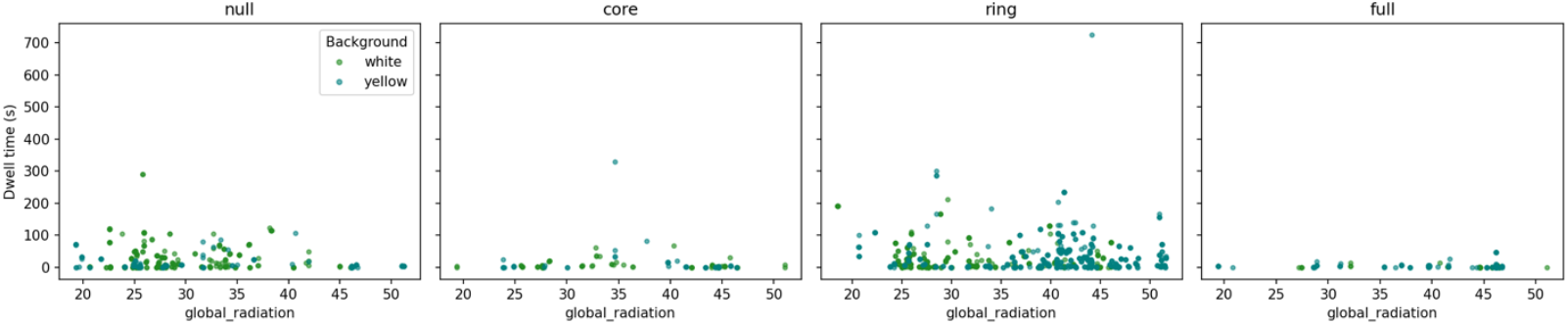
Effect of global radiation values within different UV patch allocations and artificial floral ground colour on foragers dwelling time.

From the z-standardized weather covariates, global radiation, downwelling radiation, and temperature at 2 m were all significant predictors in the foraging model (95% HDI). Global and downwelling radiation increased the odds of foraging by approximately 68% (OR ≈ 1.68) and 44% (OR ≈ 1.44), respectively, while higher temperature reduced the odds by about 34% (OR ≈ 0.66) relative to the model”s intercept. The same covariates showed consistent effects in the dwelling model (95% HDI): global and downwelling radiation prolonged dwelling time by about 1.27-fold and 1.53-fold, respectively, whereas temperature shortened it to about 0.63-fold of the baseline. Additionally, relative humidity showed marginal significance at the 90% HDI, reducing dwelling time to approximately 0.80-fold of the baseline.

The variance partitioning of the random effects revealed that in the foraging model, ∼52% of the variance was attributable to FlowerID, ∼2% to observer bias, and only ∼0.65% to bee identity, indicating strong homogeneity among individuals. The remaining ∼43% of variance was captured by the latent residual component. A similar pattern was observed for the dwelling model, with ∼31% of the variance explained by FlowerID, ∼2% by observer bias, and ∼0.75% by bee identity, while more than 64% remained in the residual. This substantial residual share suggests an underlying complexity and stochasticity in pollinator behaviour beyond the measured effects.

**Table 2.**
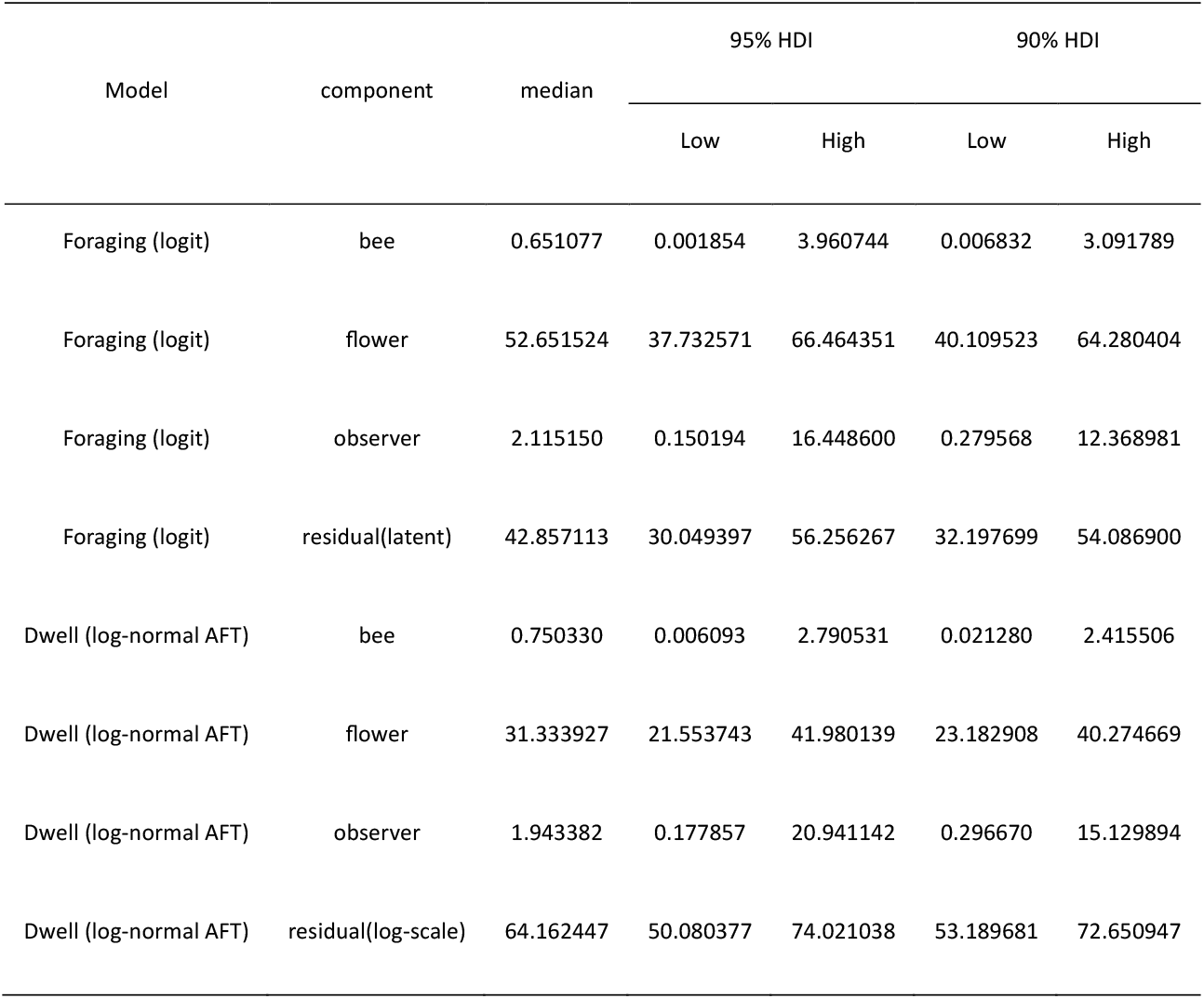
Standard deviation of noise for random effects and models reiduals at)5% HDI and 90% HDI.

Finally, when allowing for treatment-specific quadratic responses to global radiation, the estimated radiation optima differed among background colour × UV-pattern combinations, spanning approximately 25–48 J cm^−2^ across treatments. Ring-shaped UV patterns consistently showed the highest predicted foraging probabilities, with posterior mean maxima reaching p ≈ 0.87 on white backgrounds and p ≈ 0.99 on yellow backgrounds. In contrast, full UV coverage exhibited lower maxima (posterior mean p ≈ 0.40–0.74), despite overlapping radiation ranges. For several treatments, estimated optima occurred near the bounds of the observed radiation gradient, reflected by wide highest-density intervals, indicating that radiation primarily modulates the strength of visually driven preferences rather than defining a narrow absolute optimum. Detailed posterior summaries and response curves are provided in supplementary data.

## Discussion

The results of this study are broadly consistent with current knowledge on bees” foraging behaviour. The foraging model clearly reproduces the higher efficiency of sharp contrast on foraging frequency, while also hinting at the role of flower shape as a cofactor. This is evident in the repeated avoidance of artificial flowers with core UV patch allocation despite their strong contrast—contrasting with many Asterales species, where similar UV–shape combinations are effective cues. Such behaviour may further reflect the flower-loyalty of bees: in the experimental meadow, the presence of dandelions could have opposed this particular allocation by offering more appealing multimodal traits. Nevertheless, ring UV patch allocation remains the main promoter of foraging frequency/probability.

The effect of floral ground colour is another interesting outcome of this study; bees” preference for yellow over white is a well-established result of field observations, yet experiments with artificial flowers have mostly failed to capture this pattern convincingly. This duality can be traced in the handful of studies that explicitly tested colour preference with artificial flowers but ultimately shifted the discussion toward the influence of shape, symmetry, spatial placement, or contrast within the cognitive selection chain, as colour alone neither clearly reinforced nor contradicted the preference for yellow flowers (Dyer & Chittka, 2004; Giurfa et al., 1996; Lehrer et al., 1995; Orbán & Plowright, 2014). Our study successfully demonstrated the preference for yellow over white in artificial flowers, albeit positioned lower in the decision hierarchy relative to contrast. The significance levels suggest an intricate stepwise decision-making chain, which clearly accommodates contrast before floral ground colour (Randolf Menzel, 2001; Randolf Menzel & Giurfa, 2001). This statement is further supported by the dwelling model, as floral ground colour revealed no significant effect on dwelling time even at the 90% HDI, whereas contrast in the form of a ring allocation consistently led to longer foraging.

More interesting, however, is the repellent effect of full UV patch coverage observed in both models. One might argue that this phenomenon results from the loss of contrast, yet compared to the absence of any UV pattern, the full coverage effect remains evident. Interpreting this outcome requires cautious speculation, but it may be correlated with the absence of naturally occurring flowers that display full UV coverage, suggesting that bees are not evolutionarily adapted to recognize such stimuli.

Another interesting aspect of this study is how time-splines reflect bees” circadian regulated behaviour trend. The activity peak or foraging rush-hour around the noon is a well-documented pattern; in days with clear to mostly clear sky and temperature trend in range of active foraging (not optimal foraging temperature), the foraging trend advances with ascending slope, with a sharp peak around 12:00 O”clock followed by a descending curve after 13:00 O”clock. Therefore, it is no surprise that the third time-spline of this experiments which converge with this peak demonstrate a highly positive effect on both foraging frequency and its duration(Karbassioon & Stanley, 2023). A subtle yet unique phenomenon captured by our experiment is the significant dwelling time reduction at second time-spline, just prior to the foraging peak. This is a first report of a consistent dwelling trend fluctuation and very hard to interpret since more supporting data are necessary to address the ecological implication of it, as a result, we shall abstain from further elaboration on this matter.

Additional to the time-splines, the experimental day effect, representing both chronological replicas and a potential learning trend, was surprisingly insignificant. This is most likely attributable to the short duration of the experiment and the season in which it was conducted. A bee colony in summer is already fully developed and has surpassed its formative learning phase; hence, achieving a colony-wide consensus to alter an established foraging strategy within such a narrow temporal frame would be improbable. Indeed, studies have shown that while bees retain learning capacity, their behavioural plasticity declines in later season foraging contexts, favouring constancy over exploration (Baracchi, 2019; Pull et al., 2022). Notably, this absence of chronological or learning effect in our dataset is not a weakness but rather an asset, as it permits the observation of inherent cognitive responses of the focal colony, unconfounded by transient dynamic biases.

The effects of weather covariates converge well with the established biology and ethology of Apidae. As UV light is a major component of global irradiation, it naturally enhances both foraging frequency and dwelling time. Similarly, downwelling irradiation, which is strongly influenced by cloud cover, reflects sky clarity and thus higher UV input under clear conditions. By contrast, the negative effects of temperature and relative humidity likely result from late-summer conditions approaching the upper physiological boundaries for optimal foraging, with elevated temperatures directly reducing relative humidity below the desirable range(Karbassioon et al., 2023).

The standard deviations of noise further validated our experimental setup. The high variance among local replicates within each experimental day indicates that no single artificial flower overpowered others of the same ground colour and UV allocation, confirming experimental homogeneity. Observer bias was minimal, yet its presence underlines the importance of multiple assessors in studies of this kind. The negligible variation among individual bees suggests that they all belonged to a single colony and reflects the advanced developmental stage of the colony in late summer. Perhaps the most important outcome, however, is the large share of residuals in the total variance. This points to the multitude of underlying in vivo parameters and again highlights the complex nature of bees” decision-making chains. Based on the residual shares, it can be inferred that compared to foraging occurrence, the duration of foraging is determined by far more layers of information, much of which may be easily lost in background noise. Ultimately, as the decision-making chain progresses, the organism integrates the cumulative outcome of all preceding predictive steps.

By linking controlled visual traits with in vivo complexity, this study contributes to a clearer understanding of the mechanisms that shape pollination ecology and the essential role of bees in sustaining ecosystems. Finally, we hope this study will serve as a pioneer for future experiments and strongly recommend integrating additional artificial floral traits — including colour, shape, and more complex conformations — into extended experimental plots with seasonal and multi-year replicas. Furthermore, merging the two parallel models into a unified polynomial Bayesian framework would provide deeper insight into the synergistic and antagonistic interactions between foraging probability and foraging duration, offering a more holistic understanding of the bees” decision-making chain.

## Supporting information

result tables

extra graphics

python codes

## Acknowledgements

We would like to express our sincere gratitude to Dr. Christoph-Rüdiger von Bredow for his valuable suggestions, continuous support and constructive criticism that contributed to the development of our project. We also thank Dr. Dagmar Voigt and Dr. Barbara Ditsch for their guidance in the initial stages of our work and for helping us to clarify our ideas. Furthermore, we are grateful to Julien Vogelsang for his support during the implementation of our work and to Marco Matz for constructing the grid. Finally, we gratefully acknowledge the faculty of biology at TU Dresden, the Saxon State and University Library Dresden (SLUB), and Heiko Prasse from the Faculty of Meteorology for providing the necessary resources and technical equipment.

## Disclaimer

No animals, especially pollinators, were harmed during this experiment.

