## Supplementary material for "Apidae BeeHavioural response to shape-neutral visual stimuli in natural setup": result tables: summary_dwell_pub.tex

\begin{tabular}{lrrrrrrlrrrrrrrl}
\toprule
raw_term & mean & sd & hdi_3% & hdi_97% & hdi_5% & hdi_95% & term & sig95 & sig90 & sec_factor_mean & sec_factor_lo95 & sec_factor_hi95 & sec_factor_lo90 & sec_factor_hi90 & group \\
\midrule
Intercept & 1.023 & 0.537 & -0.026 & 2.058 & 0.120 & 1.928 & Intercept & False & True & 2.782 & 0.974 & 7.827 & 1.127 & 6.875 & Intercept \\
C(background, Treatment('white'))[T.yellow] & 0.087 & 0.457 & -0.792 & 0.966 & -0.655 & 0.829 & Background: yellow vs white & False & False & 1.091 & 0.453 & 2.628 & 0.519 & 2.291 & Background \\
C(uv_pattern, Treatment('null'))[T.core] & -0.023 & 0.485 & -0.979 & 0.908 & -0.838 & 0.776 & UV: core vs null & False & False & 0.977 & 0.376 & 2.480 & 0.433 & 2.172 & UV pattern \\
C(uv_pattern, Treatment('null'))[T.ring] & 0.755 & 0.460 & -0.113 & 1.680 & 0.018 & 1.505 & UV: ring vs null & False & True & 2.129 & 0.893 & 5.366 & 1.018 & 4.505 & UV pattern \\
C(uv_pattern, Treatment('null'))[T.full] & -0.897 & 0.523 & -1.904 & 0.122 & -1.751 & -0.019 & UV: full vs null & False & True & 0.408 & 0.149 & 1.129 & 0.174 & 0.981 & UV pattern \\
C(background, Treatment('white'))[T.yellow]:C(uv_pattern, Treatment('null'))[T.core] & -0.102 & 0.657 & -1.420 & 1.211 & -1.199 & 0.967 & Yellow×Core & False & False & 0.903 & 0.242 & 3.358 & 0.301 & 2.631 & Interaction \\
C(background, Treatment('white'))[T.yellow]:C(uv_pattern, Treatment('null'))[T.ring] & 0.192 & 0.599 & -0.994 & 1.350 & -0.803 & 1.174 & Yellow×Ring & False & False & 1.212 & 0.370 & 3.857 & 0.448 & 3.234 & Interaction \\
C(background, Treatment('white'))[T.yellow]:C(uv_pattern, Treatment('null'))[T.full] & 0.742 & 0.648 & -0.501 & 2.008 & -0.300 & 1.805 & Yellow×Full & False & False & 2.101 & 0.606 & 7.448 & 0.741 & 6.078 & Interaction \\
time_spline_3 & 0.124 & 0.527 & -0.926 & 1.129 & -0.764 & 0.982 & time_spline_3 & False & False & 1.132 & 0.396 & 3.093 & 0.466 & 2.669 & Time/Day \\
time_spline_2 & 1.125 & 0.550 & 0.051 & 2.171 & 0.220 & 2.007 & time_spline_2 & True & True & 3.080 & 1.053 & 8.771 & 1.246 & 7.442 & Time/Day \\
day_num_z & -0.234 & 0.222 & -0.666 & 0.200 & -0.593 & 0.127 & Learning (day, z) & False & False & 0.791 & 0.514 & 1.221 & 0.553 & 1.135 & Time/Day \\
time_spline_0 & 0.049 & 0.634 & -1.198 & 1.266 & -0.994 & 1.091 & time_spline_0 & False & False & 1.050 & 0.302 & 3.548 & 0.370 & 2.977 & Time/Day \\
time_spline_1 & -1.234 & 0.605 & -2.430 & -0.096 & -2.220 & -0.265 & time_spline_1 & True & True & 0.291 & 0.088 & 0.909 & 0.109 & 0.767 & Time/Day \\
z_global_radiation & 0.236 & 0.121 & -0.001 & 0.471 & 0.038 & 0.432 & Global rad (z) & False & True & 1.266 & 0.999 & 1.601 & 1.039 & 1.540 & Weather \\
z_temp_2m & -0.473 & 0.219 & -0.909 & -0.055 & -0.842 & -0.119 & Temp (z) & True & True & 0.623 & 0.403 & 0.947 & 0.431 & 0.888 & Weather \\
z_wind_avg & -0.027 & 0.069 & -0.159 & 0.107 & -0.139 & 0.086 & Wind (z) & False & False & 0.973 & 0.853 & 1.113 & 0.870 & 1.090 & Weather \\
z_humidity_rel & -0.221 & 0.134 & -0.485 & 0.050 & -0.441 & -0.001 & Humidity (z) & False & True & 0.802 & 0.616 & 1.051 & 0.643 & 0.999 & Weather \\
z_downwelling_radiation & 0.426 & 0.086 & 0.262 & 0.593 & 0.287 & 0.570 & Downwelling rad (z) & True & True & 1.531 & 1.299 & 1.810 & 1.333 & 1.768 & Weather \\
\bottomrule
\end{tabular}
