## Supplementary material for "Apidae BeeHavioural response to shape-neutral visual stimuli in natural setup": result tables: summary_foraging_pub.tex

\begin{tabular}{lrrrrrrlrrrrrrrrrrrrl}
\toprule
raw_term & mean & sd & hdi_3% & hdi_97% & hdi_5% & hdi_95% & term & sig95 & sig90 & odds_ratio_mean & odds_ratio_lo95 & odds_ratio_hi95 & odds_ratio_lo90 & odds_ratio_hi90 & p(baseline+beta)_mean & p(baseline+beta)_lo95 & p(baseline+beta)_hi95 & p(baseline+beta)_lo90 & p(baseline+beta)_hi90 & group \\
\midrule
Intercept & -0.631 & 0.707 & -2.036 & 0.726 & -1.799 & 0.514 & Intercept & False & False & 0.532 & 0.131 & 2.067 & 0.165 & 1.671 & 0.221 & 0.065 & 0.524 & 0.081 & 0.471 & Intercept \\
C(background, Treatment('white'))[T.yellow] & 1.264 & 0.677 & -0.078 & 2.594 & 0.161 & 2.395 & Background: yellow vs white & False & True & 3.538 & 0.925 & 13.385 & 1.174 & 10.970 & 0.653 & 0.330 & 0.877 & 0.385 & 0.854 & Background \\
C(uv_pattern, Treatment('null'))[T.core] & -0.273 & 0.758 & -1.766 & 1.214 & -1.539 & 0.956 & UV: core vs null & False & False & 0.761 & 0.171 & 3.367 & 0.215 & 2.601 & 0.288 & 0.083 & 0.642 & 0.103 & 0.581 & UV pattern \\
C(uv_pattern, Treatment('null'))[T.ring] & 0.421 & 0.715 & -0.933 & 1.806 & -0.730 & 1.609 & UV: ring vs null & False & False & 1.523 & 0.393 & 6.084 & 0.482 & 4.999 & 0.448 & 0.173 & 0.764 & 0.204 & 0.727 & UV pattern \\
C(uv_pattern, Treatment('null'))[T.full] & -1.288 & 0.792 & -2.873 & 0.263 & -2.587 & 0.006 & UV: full vs null & False & False & 0.276 & 0.057 & 1.301 & 0.075 & 1.006 & 0.128 & 0.029 & 0.409 & 0.038 & 0.349 & UV pattern \\
C(background, Treatment('white'))[T.yellow]:C(uv_pattern, Treatment('null'))[T.core] & -0.416 & 0.946 & -2.226 & 1.482 & -1.958 & 1.162 & Yellow×Core & False & False & 0.660 & 0.108 & 4.404 & 0.141 & 3.196 & 0.260 & 0.054 & 0.701 & 0.070 & 0.630 & Interaction \\
C(background, Treatment('white'))[T.yellow]:C(uv_pattern, Treatment('null'))[T.ring] & 0.191 & 0.889 & -1.548 & 1.968 & -1.222 & 1.683 & Yellow×Ring & False & False & 1.211 & 0.213 & 7.157 & 0.295 & 5.384 & 0.392 & 0.102 & 0.792 & 0.136 & 0.741 & Interaction \\
C(background, Treatment('white'))[T.yellow]:C(uv_pattern, Treatment('null'))[T.full] & 0.807 & 0.952 & -1.042 & 2.690 & -0.770 & 2.373 & Yellow×Full & False & False & 2.241 & 0.353 & 14.727 & 0.463 & 10.735 & 0.544 & 0.158 & 0.887 & 0.198 & 0.851 & Interaction \\
time_spline_3 & 0.589 & 0.820 & -0.987 & 2.229 & -0.752 & 1.950 & time_spline_3 & False & False & 1.801 & 0.373 & 9.294 & 0.471 & 7.030 & 0.489 & 0.166 & 0.832 & 0.201 & 0.789 & Time/Day \\
time_spline_2 & 2.571 & 0.782 & 1.050 & 4.158 & 1.285 & 3.841 & time_spline_2 & True & True & 13.077 & 2.858 & 63.974 & 3.616 & 46.582 & 0.874 & 0.603 & 0.971 & 0.658 & 0.961 & Time/Day \\
day_num_z & -0.235 & 0.383 & -0.977 & 0.503 & -0.854 & 0.399 & Learning (day, z) & False & False & 0.791 & 0.376 & 1.653 & 0.426 & 1.490 & 0.296 & 0.167 & 0.468 & 0.185 & 0.442 & Time/Day \\
time_spline_0 & 0.586 & 0.893 & -1.179 & 2.294 & -0.891 & 2.039 & time_spline_0 & False & False & 1.797 & 0.308 & 9.917 & 0.410 & 7.684 & 0.489 & 0.141 & 0.841 & 0.179 & 0.804 & Time/Day \\
time_spline_1 & -0.963 & 0.897 & -2.666 & 0.801 & -2.417 & 0.547 & time_spline_1 & False & False & 0.382 & 0.070 & 2.227 & 0.089 & 1.728 & 0.169 & 0.036 & 0.542 & 0.045 & 0.479 & Time/Day \\
z_global_radiation & 0.522 & 0.213 & 0.099 & 0.944 & 0.167 & 0.874 & Global rad (z) & True & True & 1.686 & 1.104 & 2.569 & 1.182 & 2.396 & 0.473 & 0.370 & 0.578 & 0.386 & 0.561 & Weather \\
z_temp_2m & -1.081 & 0.376 & -1.819 & -0.353 & -1.708 & -0.476 & Temp (z) & True & True & 0.339 & 0.162 & 0.703 & 0.181 & 0.621 & 0.153 & 0.079 & 0.272 & 0.088 & 0.248 & Weather \\
z_wind_avg & -0.208 & 0.134 & -0.472 & 0.058 & -0.428 & 0.007 & Wind (z) & False & False & 0.813 & 0.624 & 1.060 & 0.652 & 1.007 & 0.302 & 0.249 & 0.361 & 0.257 & 0.349 & Weather \\
z_humidity_rel & -0.328 & 0.256 & -0.846 & 0.174 & -0.760 & 0.085 & Humidity (z) & False & False & 0.720 & 0.429 & 1.191 & 0.468 & 1.088 & 0.277 & 0.186 & 0.388 & 0.199 & 0.367 & Weather \\
z_downwelling_radiation & 0.362 & 0.154 & 0.064 & 0.665 & 0.113 & 0.618 & Downwelling rad (z) & True & True & 1.437 & 1.066 & 1.944 & 1.120 & 1.854 & 0.433 & 0.362 & 0.508 & 0.374 & 0.497 & Weather \\
\bottomrule
\end{tabular}
