## Supplementary material for "Apidae BeeHavioural response to shape-neutral visual stimuli in natural setup": extra graphics: Extra Graphics.pdf

### Appendix

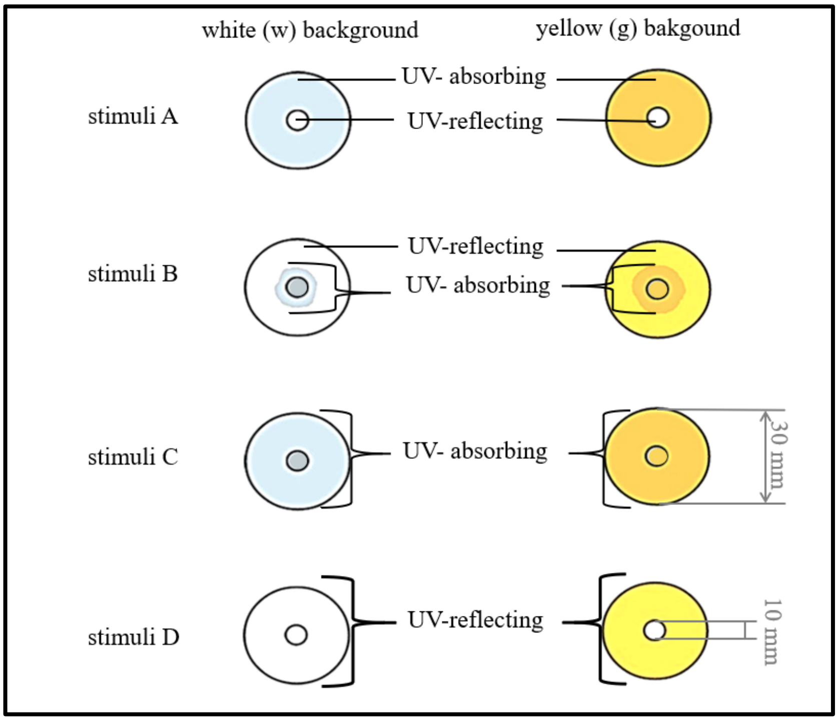


Figure 1: Design of the white and yellow stimuli. Stimulus A: UV-absorbing outer part, UV-reflecting inner part; stimulus B: UV-reflecting outer part, UV-absorbing inner part; stimulus C: UV-absorbing outer and inner part; stimulus D: UV-reflecting outer and inner part.

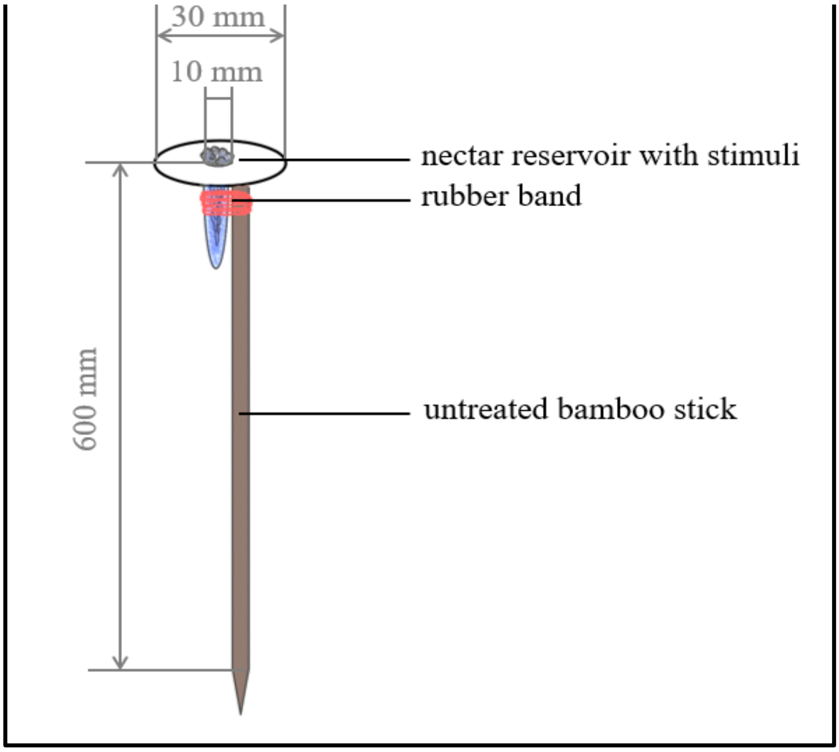


Figure 2: Schematic diagram of fully assembled artificial flower: A centrifuge tube filled with cotton wool and nectar solution was inserted through the hole of the paper ring and attached to an untreated bamboo stick using a rubber band.

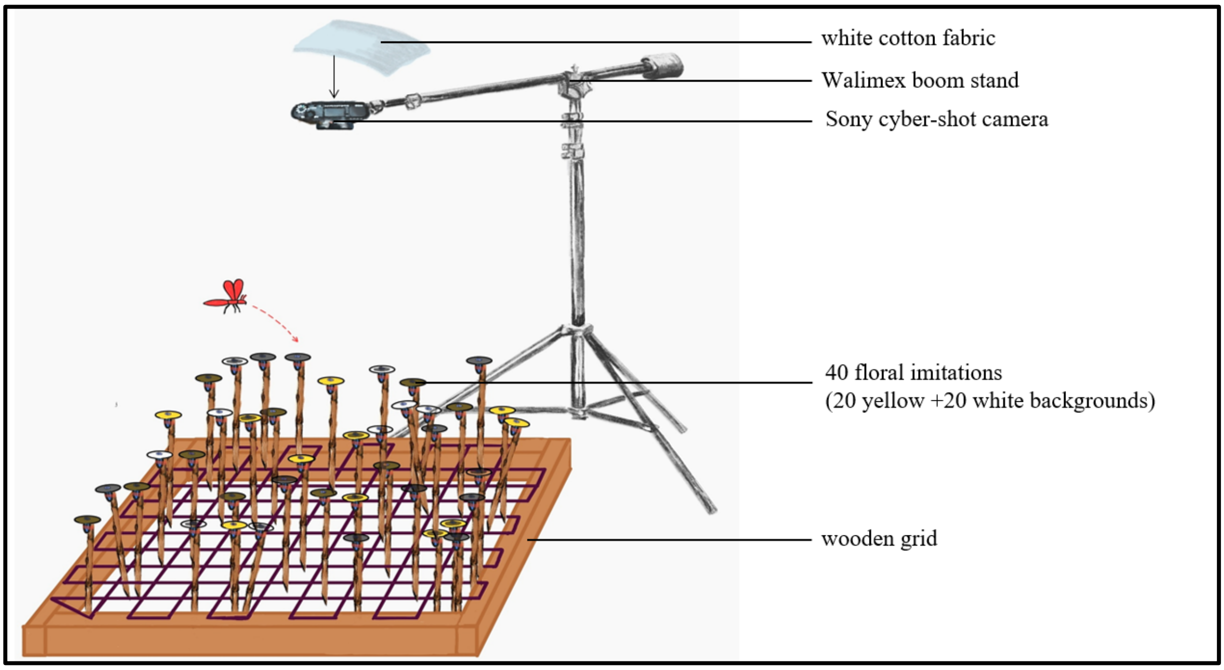


Figure 3: Schematic sketch of the experimental setup. The stimuli were distributed in the randomly placed ecological sample quadrat. Above the quadrat a camera was installed using a boom stand, where all stimuli could be seen.

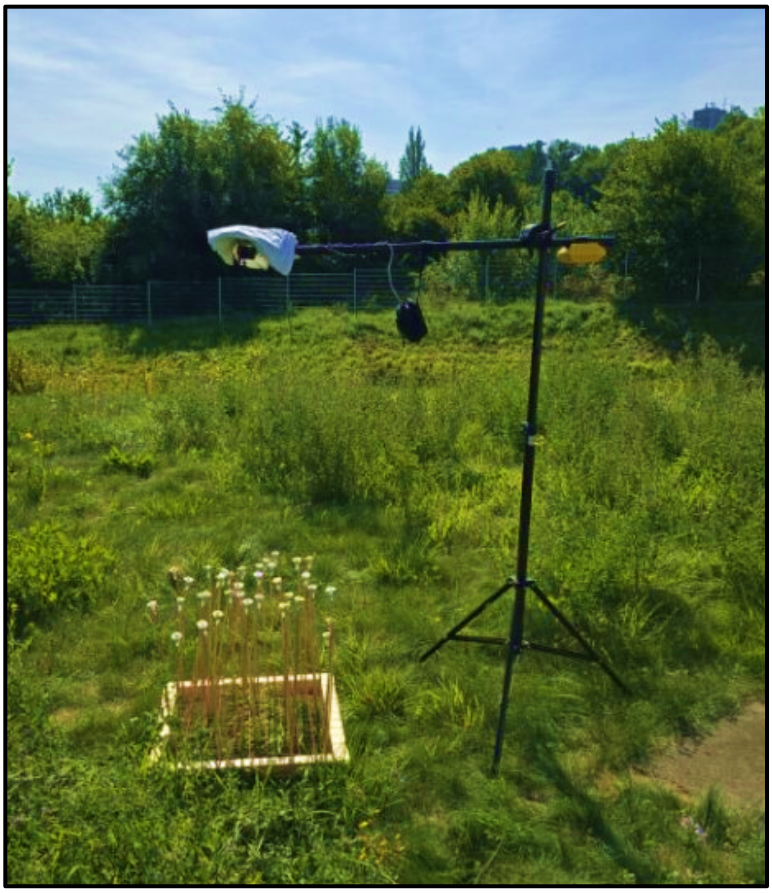

Figure 4: Experimental setup in the field. The stimuli were distributed in the randomly placed ecological sample quadrat. Above the quadrat a camera was installed using a boom stand, where all stimuli could be seen.
