## Supplementary figures and images for "Apidae BeeHavioural response to shape-neutral visual stimuli in natural setup"

### dwell_hist.png

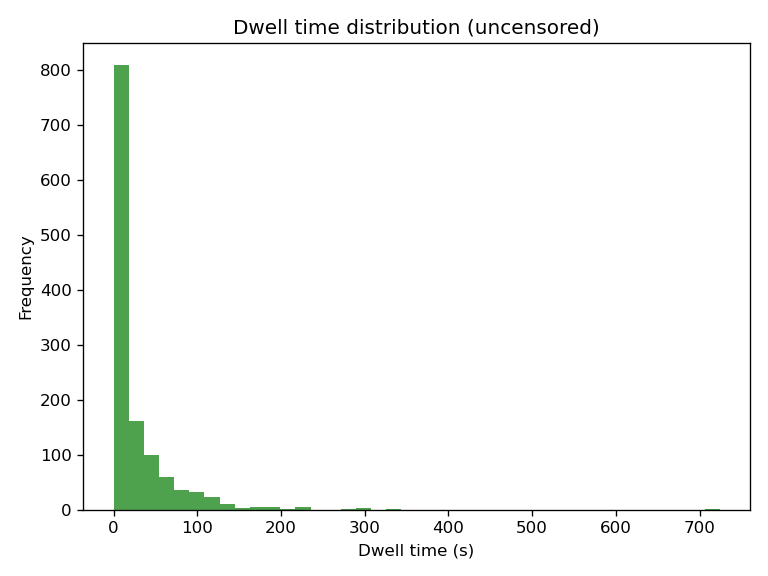

### forest_dwell.png

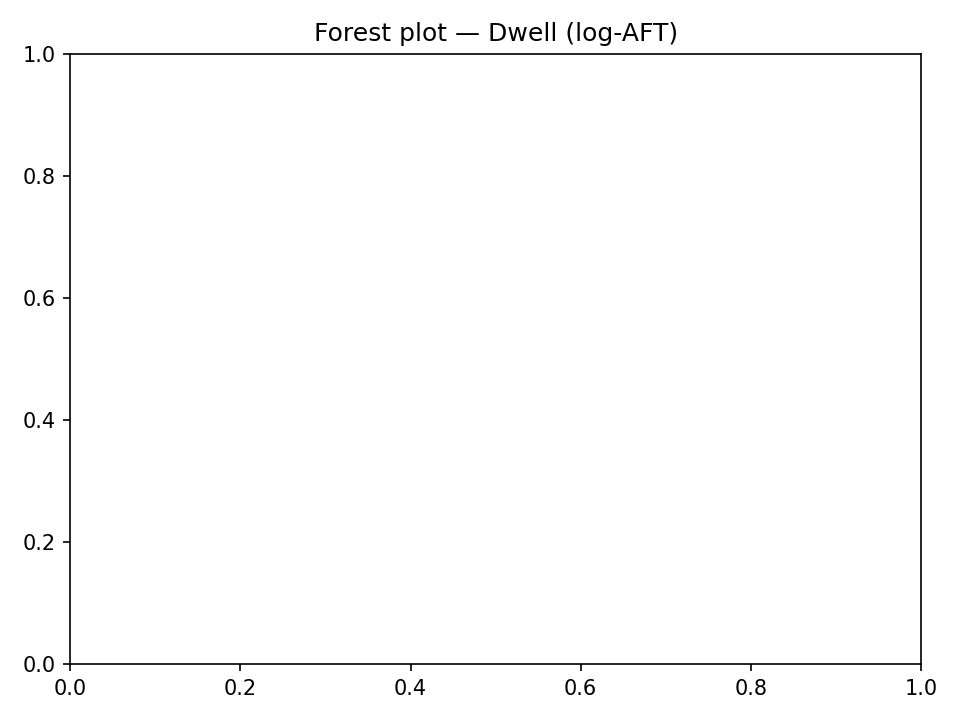

### forest_dwell_custom.png

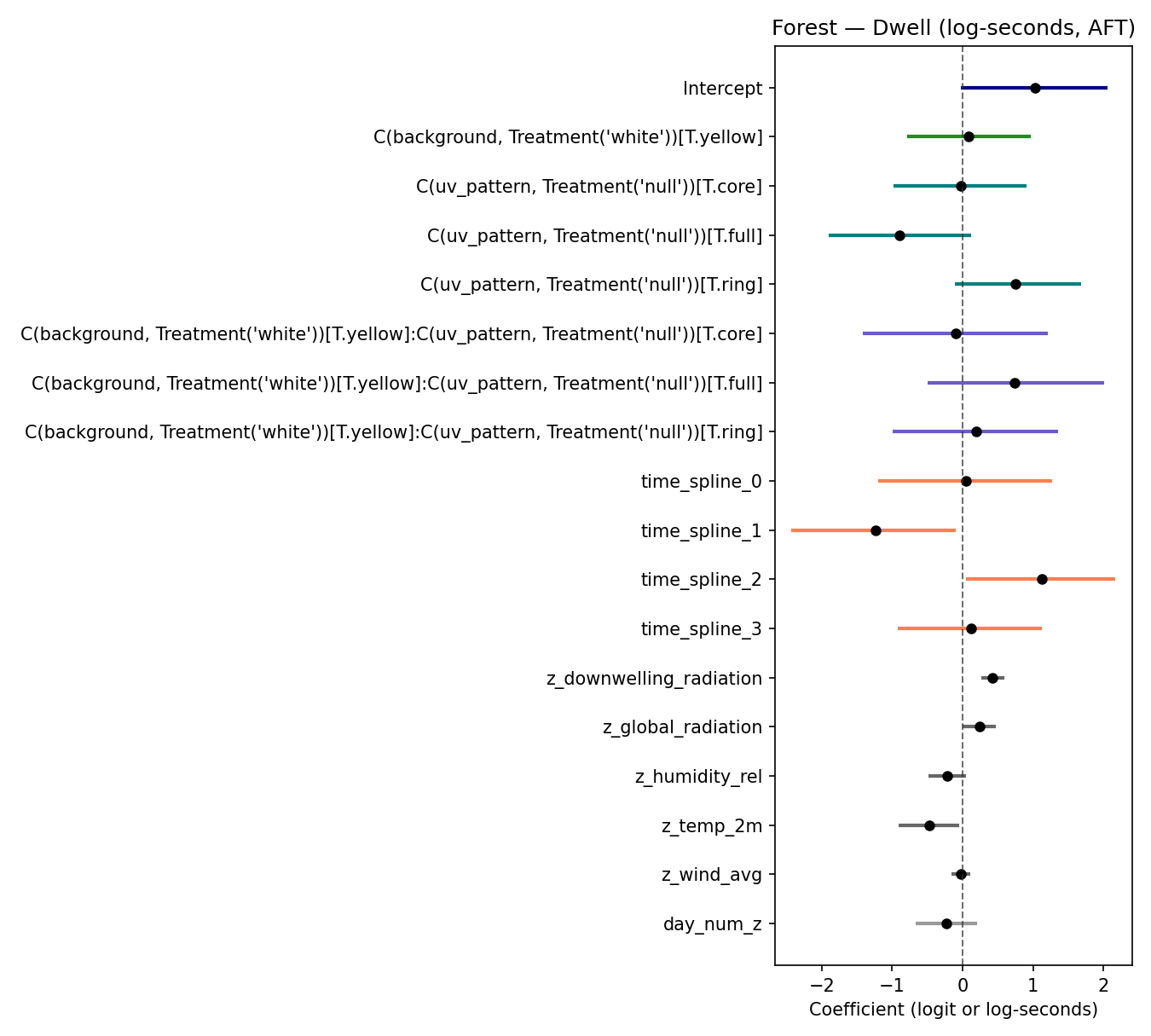

### forest_foraging.png

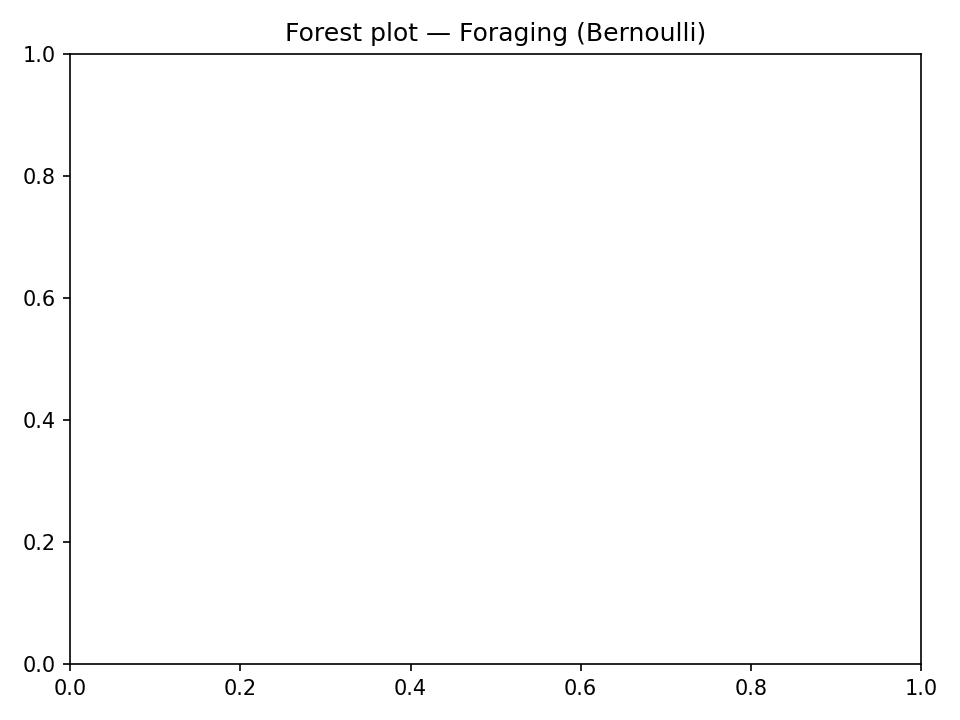

### forest_foraging_custom.png

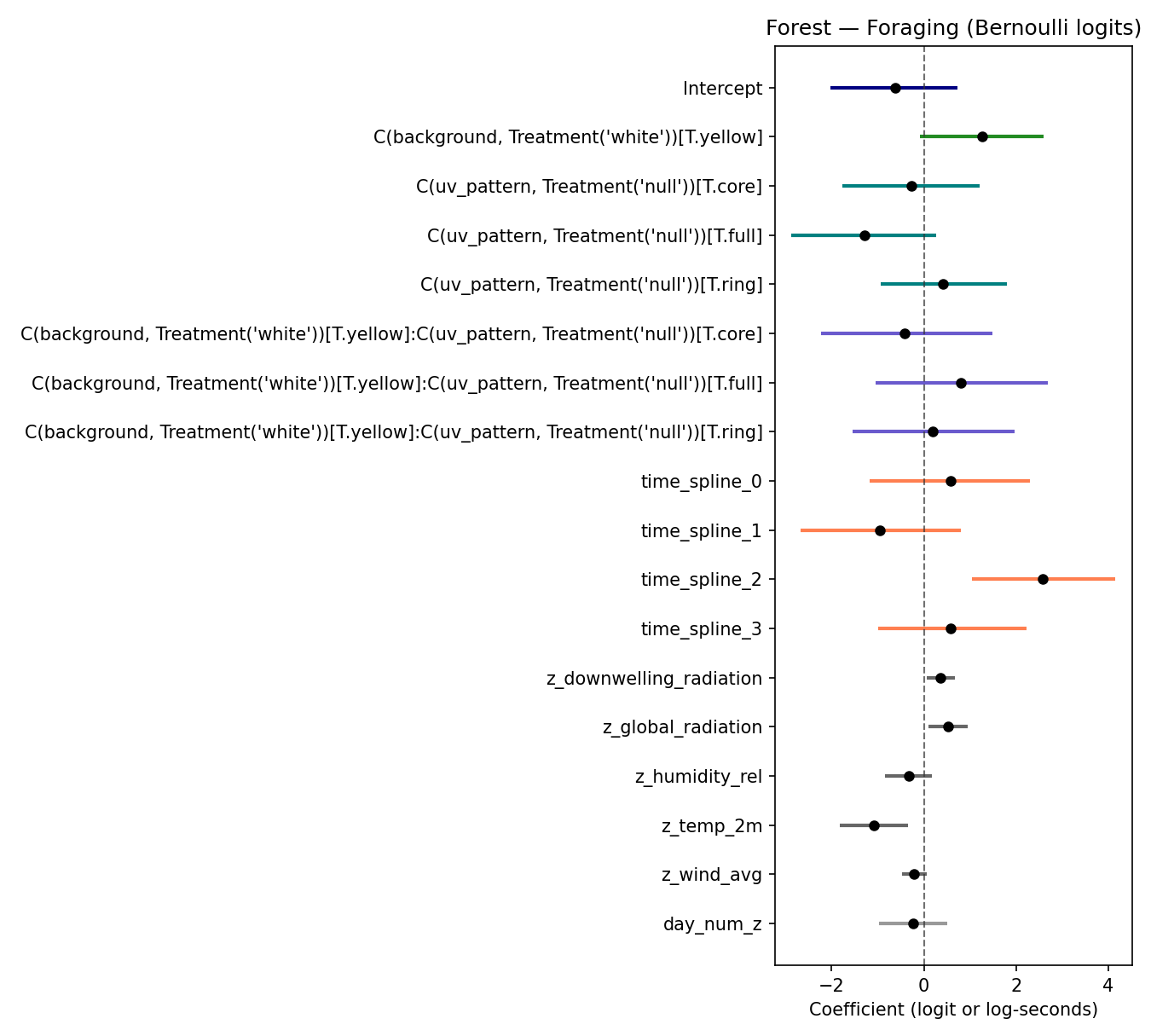

### learning_curve.png

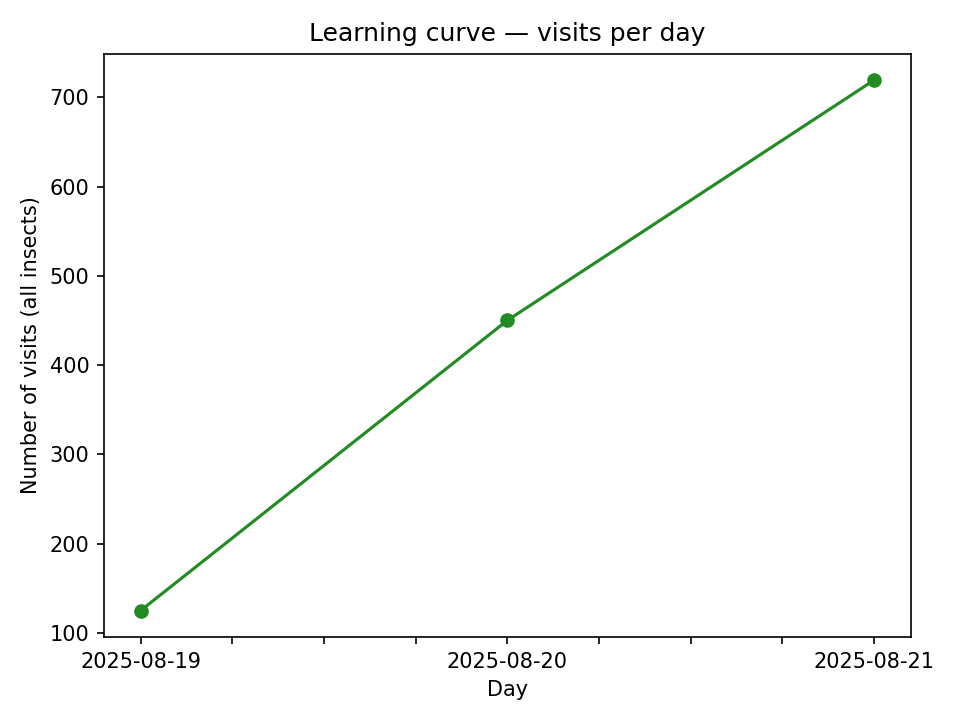

### ppc_dwell_log_overlay.png

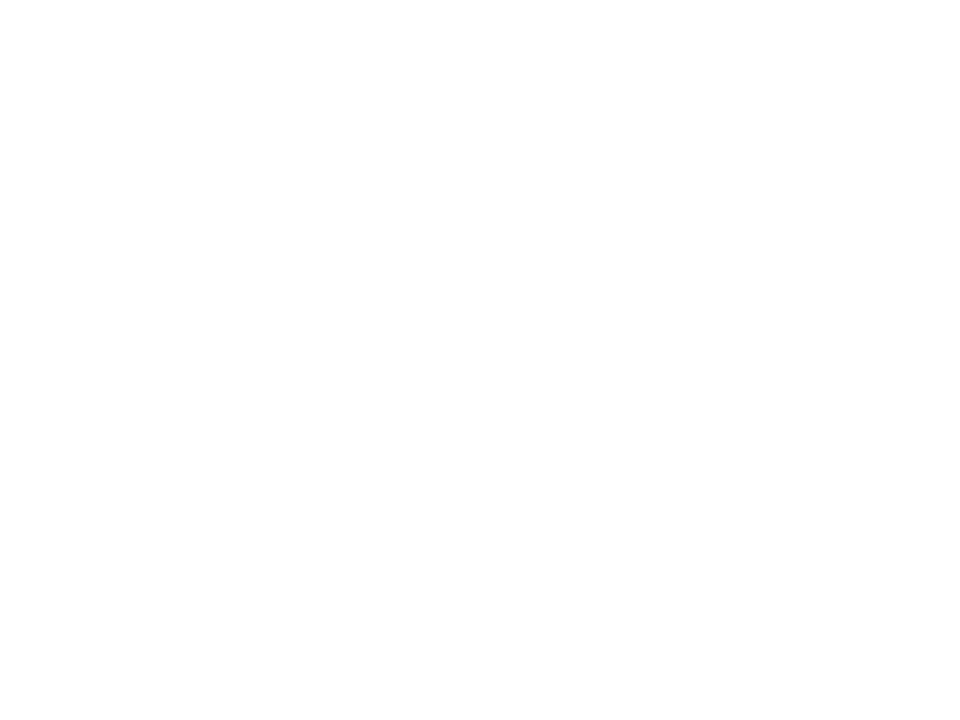

### ppc_dwell_seconds_unc.png

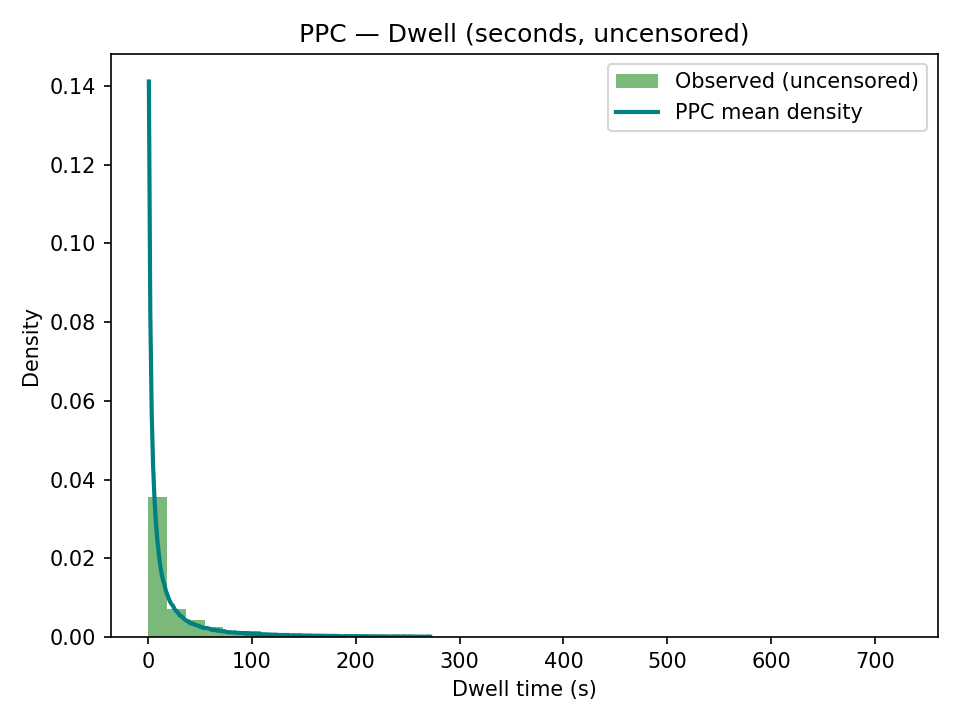

### ppc_foraging_calibration.png

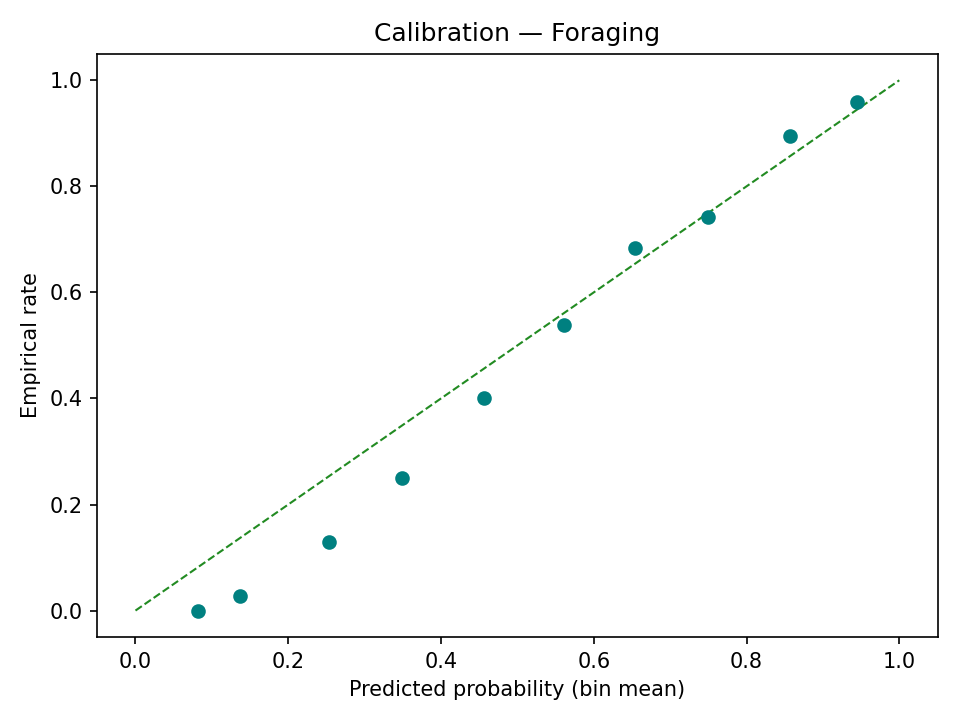

### qc_downrad_vs_time.png

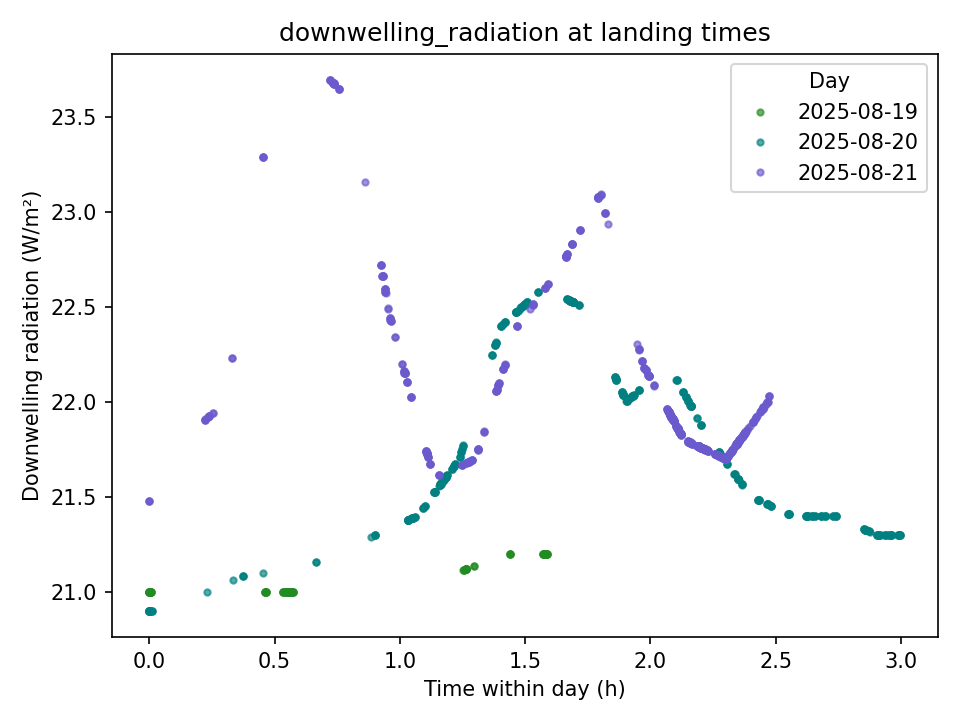

### qc_globalrad_vs_time.png

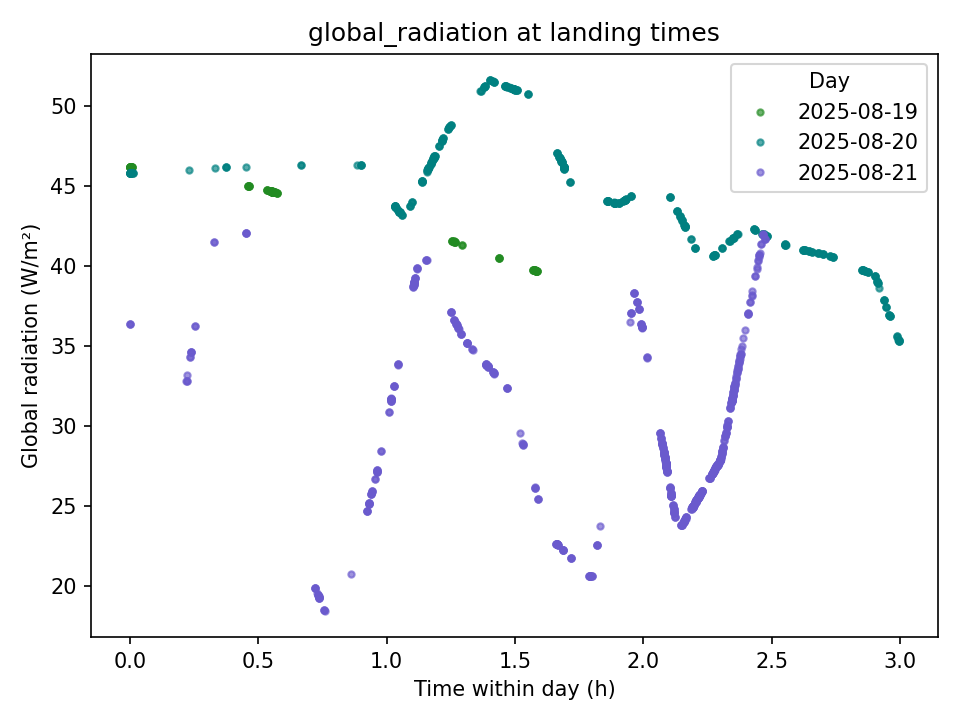

### qc_hum_vs_time.png

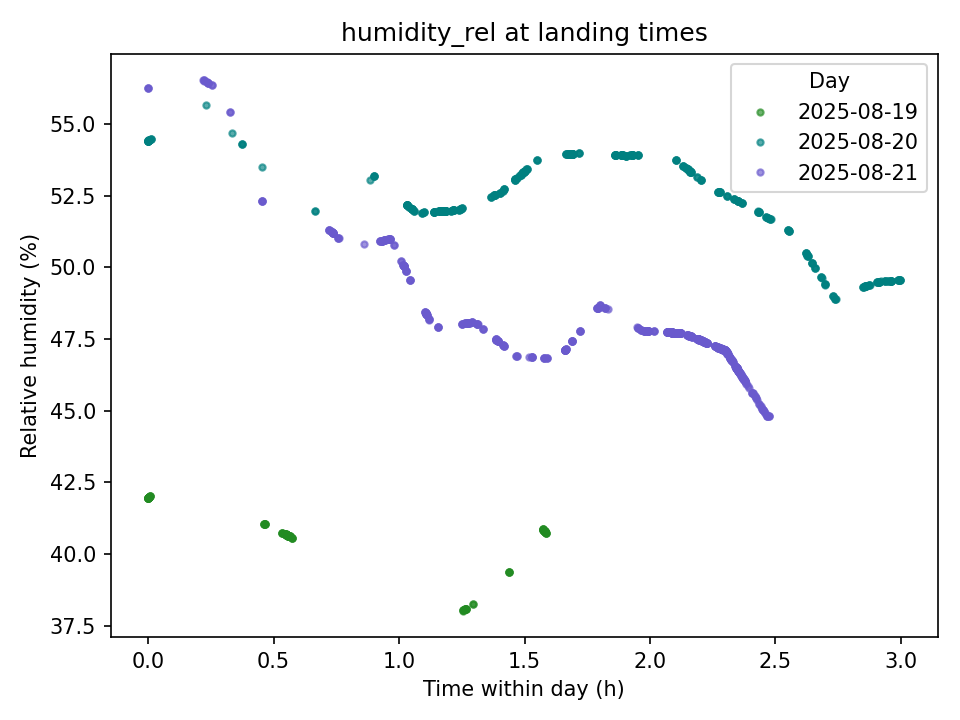

### qc_scatter_dwell_vs_global.png

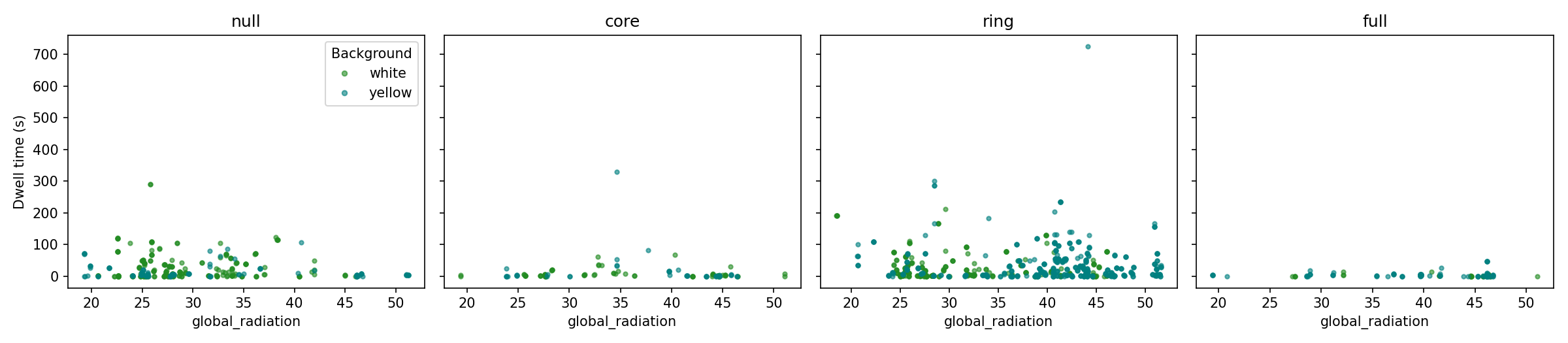

### qc_scatter_dwell_vs_wind.png

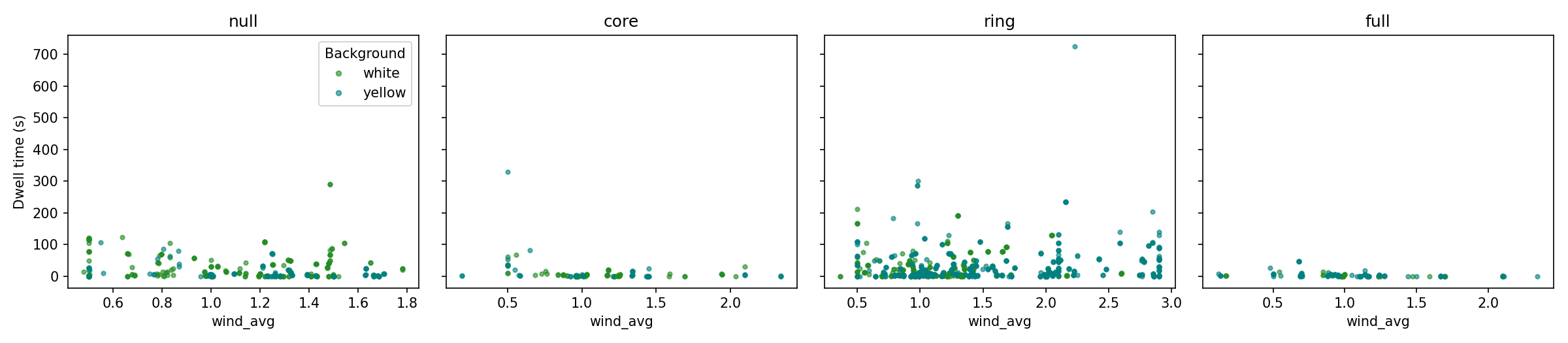

### qc_temp_vs_time.png

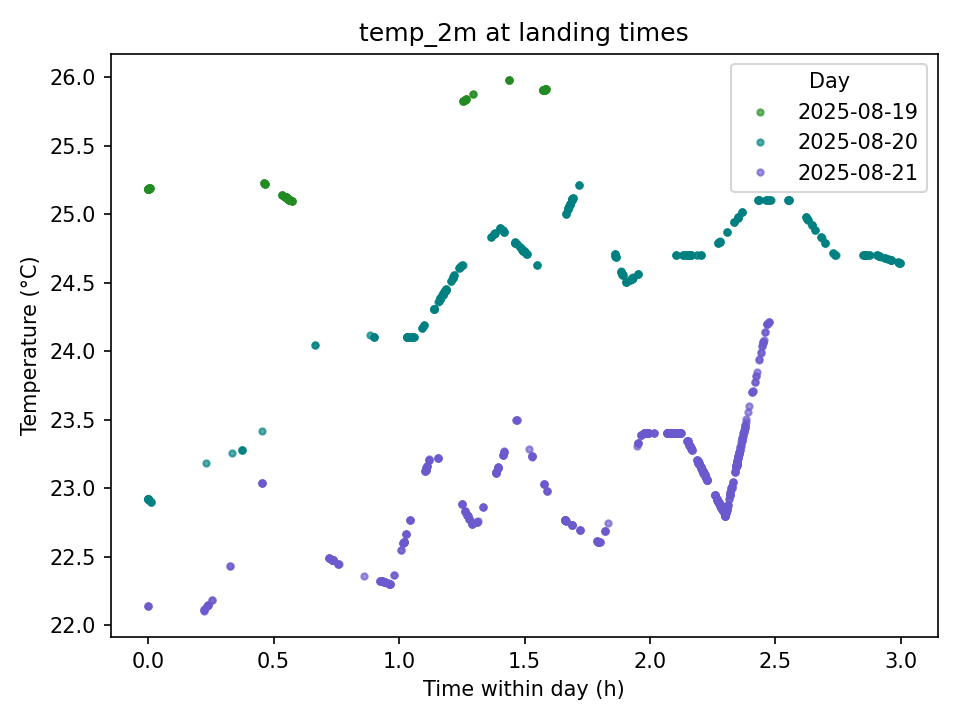

### qc_wind_vs_time.png

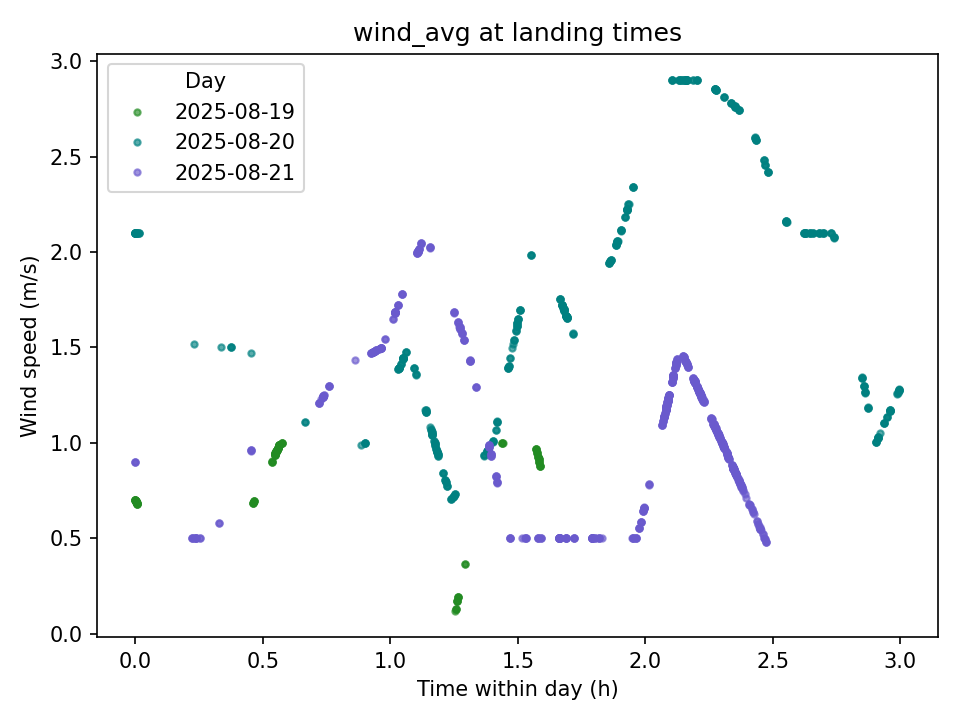

### timecurve_dwell.png

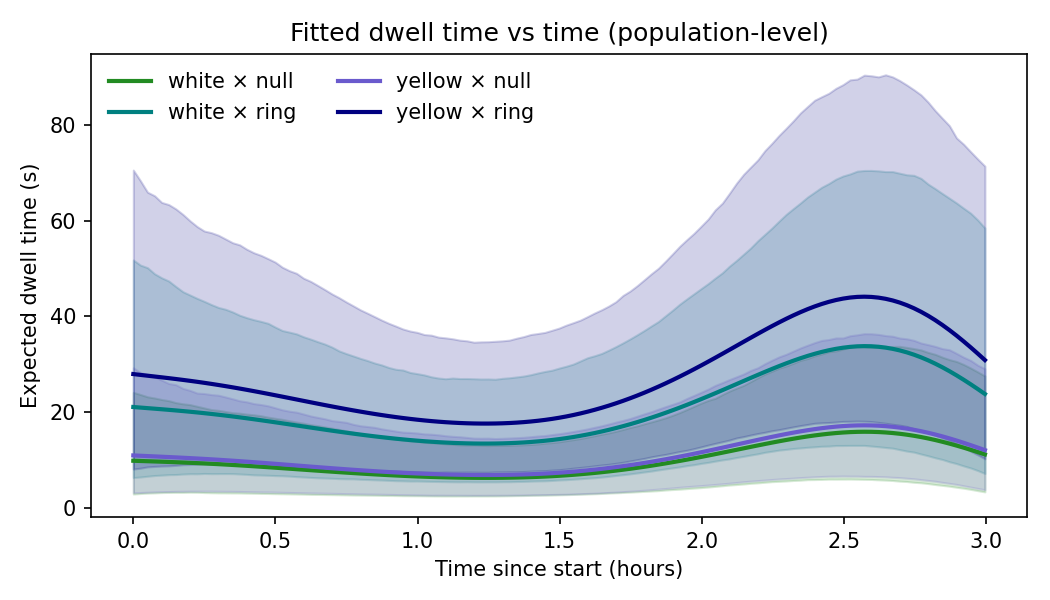

### timecurve_foraging.png

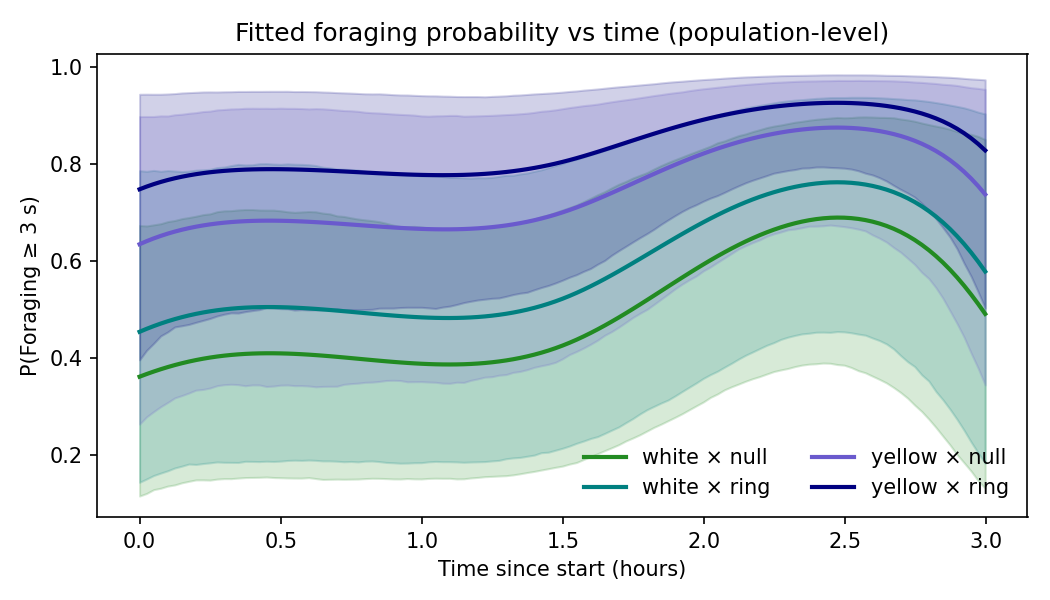
