## Supplementary material for "Apidae BeeHavioural response to shape-neutral visual stimuli in natural setup": python codes: BeeHavior_2.docx

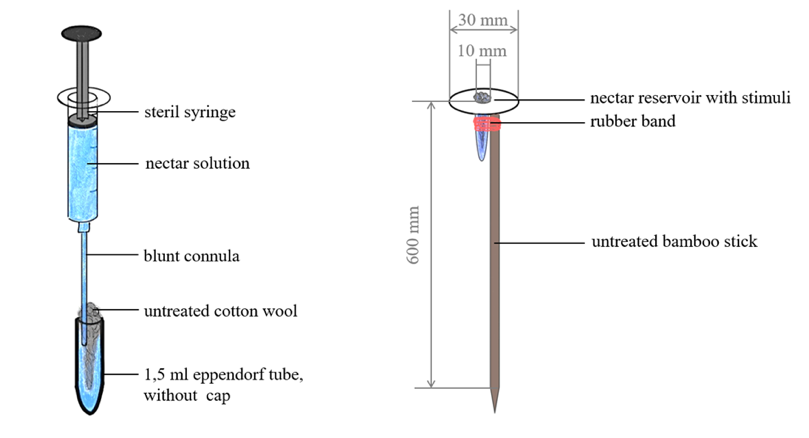

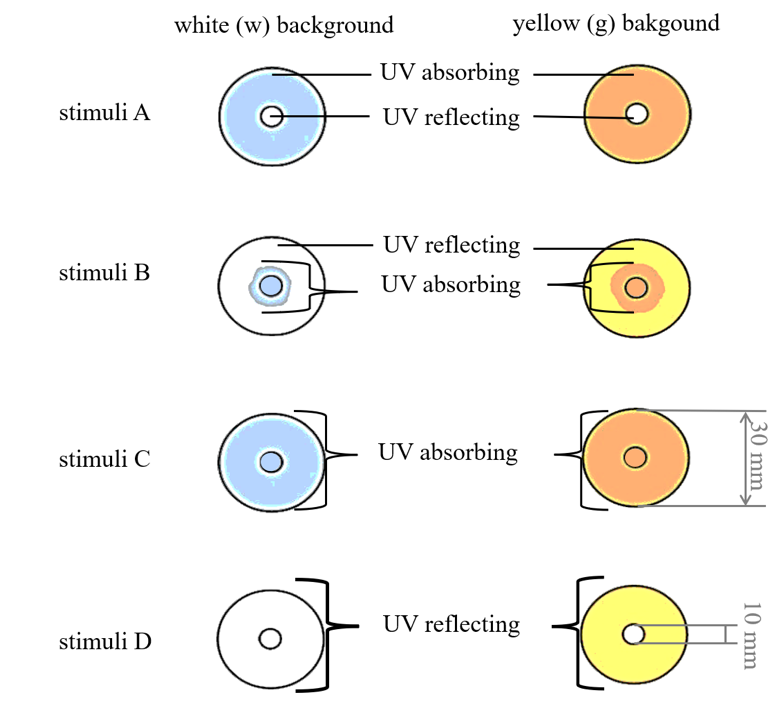
The UV patterns were applied daily prior to the experiment by spraying the designated sections with a commercially available transparent sunscreen spray (Eucerin® Sun Protection, 50+ SPF, Oil Control; active substances: Diethylamino Hydroxybenzoyl Hexyl Benzoate, Ethylhexyl Salicylate). For stimulus A (ring), the entire paper ring was treated; for stimulus B (core), only the dome-shaped cotton wick in the center was sprayed; for stimulus C (full), both the paper ring and the central wick were treated, while stimulus D (null) remained untreated.The pattern application was validated using UV imaging with a Sony Alpha NEX-5T mirrorless camera (Quartz sensor window) equipped with a Sigma 30 mm f/2.8 DN Art lens, a No. 2 Kenko close-up filter, and a Baader Venus U-filter.

Figure 1 Design of the white and yellow stimuli.

The assembled artificial flowers were mounted on approximately 60 cm long bamboo sticks (ø ≥ 0.7 cm) using rubber bands. Reservoirs were filled to the 1.0 ml mark with freshly thawed aliquots of artificial nectar (prepared on 18 August and stored at −20 °C), consisting of 30% (w/v) sucrose, 0.6% (v/v) linalool, and 0.1% (v/v) phenylacetaldehyde.

Figure 2 Schematic diagram of fully assembled artificial flower

Plot randomization was achieved by throwing an ecological sampling quadrat (50 × 50 cm, subdivided into 100 cells of 5 × 5 cm). Vegetation within the selected quadrat was flattened to reduce competition. The 40 stimuli were then distributed under the following constraints: (i) no more than 20 cells contained a single stimulus, (ii) cells could contain two stimuli of either the same or different colours, (iii) up to three stimuli were allowed per cell if they were either all of the same colour or a combination of two of one colour plus one of another, and (iv) within any 3 × 3 block of cells, no more than six stimuli of the same colour were permitted.

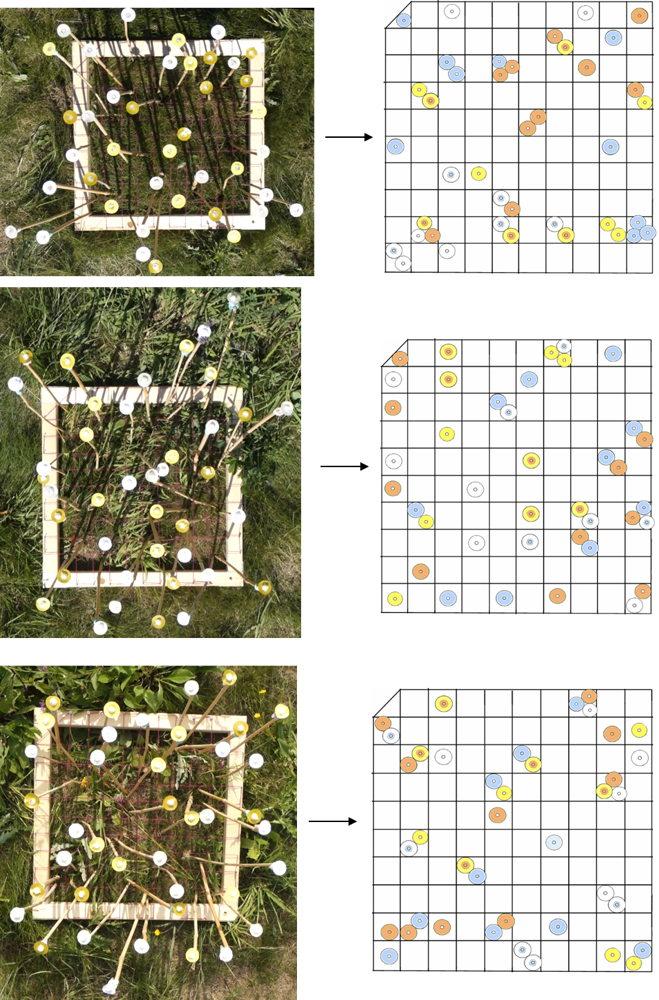
A Sony Cyber-shot DSC-RX100M3 camera was mounted horizontally above the experimental plot on a Wallimex WT-501 boom stand, and continuous 3-hour recordings of pollinator interactions with artificial flowers were made between 10:00 and 14:00.

Figure 3 Map of artificial distribution within each experimental day (day 1 to 3 from above)

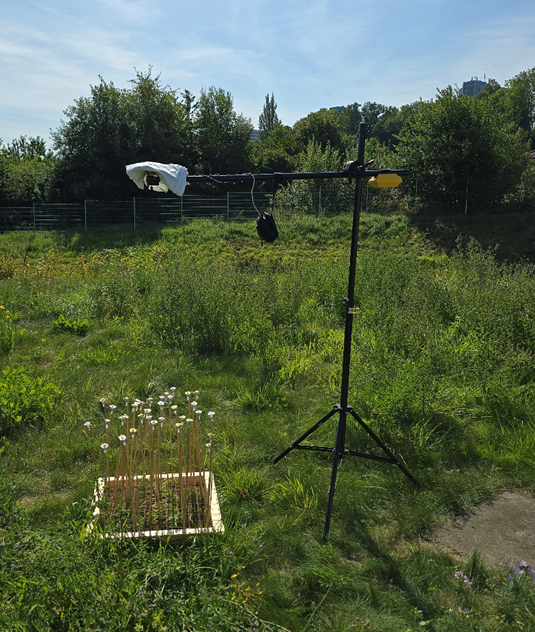

Figure 4 A view of experimental setup during recording

All video recordings from the three experimental days were surveyed independently by four observers. Observers used a time-stamped, event-based evaluation sheet to document pollinator activity. For each visit, the observer assigned a Visitor identity (VisitorID) to the individual pollinator and recorded the event type (landing or take-off) on a given artificial flower (FlowerID). To ensure consistency, each VisitorID was retained for as long as the same individual remained within the recording frame, and reassigned only when a pollinator left the frame and re-entered. The family and genus of the visitor were noted in the Category column. Time-stamps were extracted by Shotcut software (Meltytech, LLC, 2025). Each observer completed the full survey of all three experimental days individually, ensuring complete replication of annotations across observers.

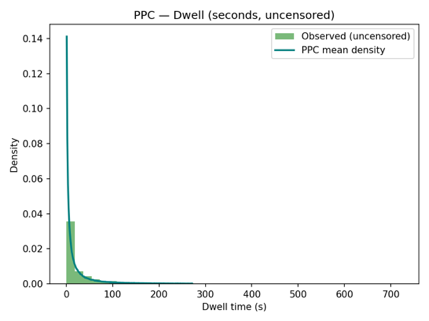

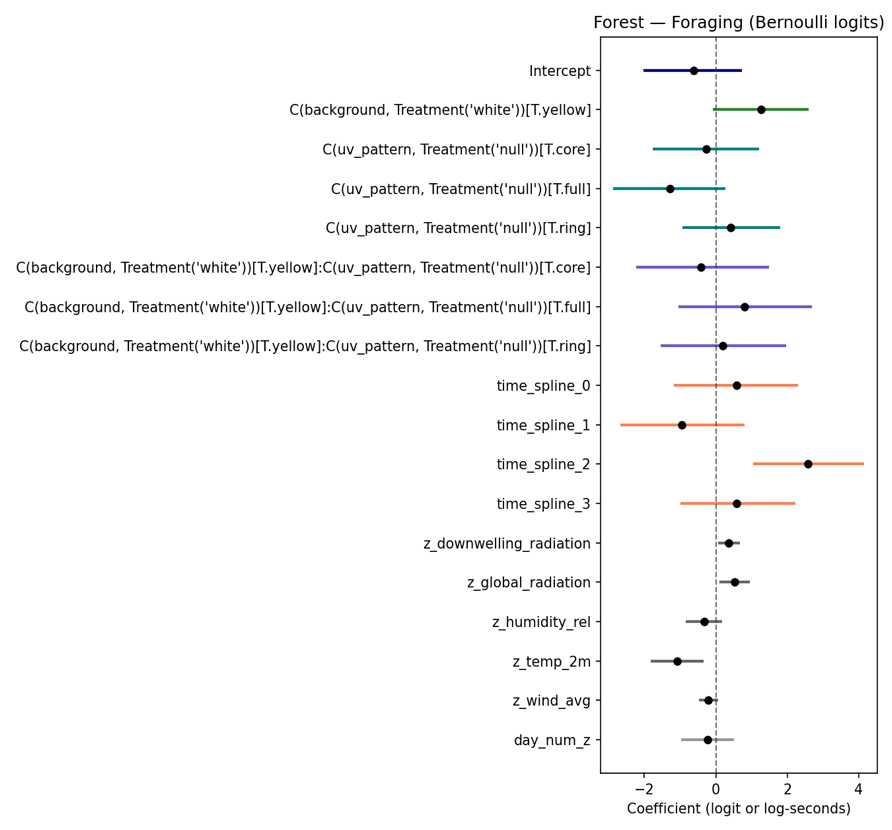
Results:
Both the foraging (Bernoulli–logit) and dwelling-time (log-normal AFT) models demonstrated excellent convergence and fit. Across all parameters, the Gelman–Rubin statistic ( r̂ ) was equal to 1.00, effective sample sizes (ESS) exceeded 1,000 for both bulk and tail estimates, and no divergent transitions were reported. Posterior predictive checks indicated strong agreement between observed and model-simulated data, supporting the adequacy of model specification.

Figure 6 PPC of dwelling time (above) and foraging (below) model, revealing a very good agreement between posterior simulated and observed data

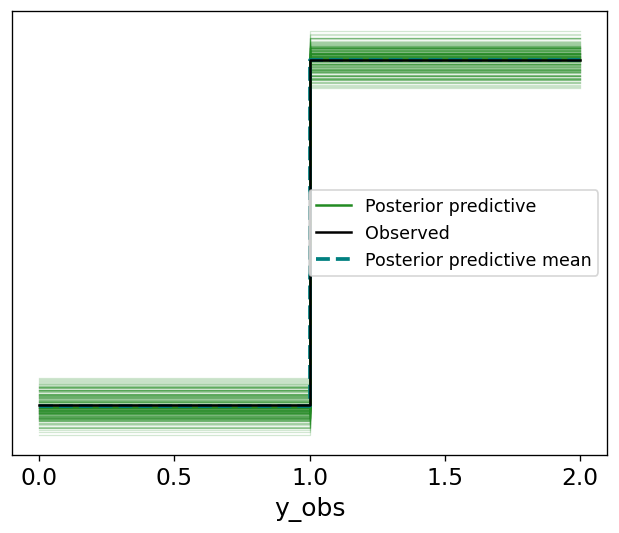

The Bernoulli/logit foraging model showed no effects of background colour or UV allocation at the 95% HDI. At the 90% HDI, however, flowers with a yellow background were 27% more likely to be foraged than those with a white background, and flowers with a full UV pattern were 18.5% less likely to be foraged than those without UV.

| contrast | Log Odds 95% HDI | | | Log Odds 90% HDI | | | Odds Ratio 95% HDI | | Odds Ratio 90% HDI | |
| --- | --- | --- | --- | --- | --- | --- | --- | --- | --- | --- |
|  | Low | High | Sig. | Low | High | Sig. | Low | High | Low | High |
| Foraging: white (ring − full) | -0.07 | 3.52 | x | 0.21 | 3.22 | * | 0.93 | 33.81 | 1.23 | 25.06 |
| Foraging: yellow (ring − full) | -0.65 | 2.80 | x | -0.37 | 2.54 | x | 0.52 | 16.42 | 0.69 | 12.71 |
| Foraging: UV=null  (yellow − white) | -0.08 | 2.60 | x | 0.16 | 2.40 | * | 0.92 | 13.39 | 1.17 | 10.97 |
| Foraging: white (ring − null) | -0.93 | 1.81 | x | -0.73 | 1.61 | x | 0.39 | 6.08 | 0.48 | 4.10 |
| Foraging: yellow (ring − null) | -1.03 | 2.26 | x | -0.80 | 1.97 | x | 0.36 | 9.55 | 0.45 | 7.14 |

Table 1 Comparison between log odds of foraging within different fixed effect combination at 95% HDI and 90% HDI

In the AFT/log-normal model, the intercept was significant at the 95% HDI, indicating that even under baseline conditions (white background, no UV pattern, average weather), dwell times were systematically longer than expected under a neutral model. Specifically, baseline foraging events were estimated to last up to ~2.8 times longer, highlighting an inherent structure in dwelling behaviour independent of treatment effects. Neither background colour nor UV pattern allocation showed significant effects at the 95% HDI. However, at the more permissive 90% HDI, dwell times were likely ~2.1 times longer on flowers with ring-shaped UV allocation, while flowers with full UV coverage were associated with ~0.6 times shorter dwell times compared to no UV pattern at all.

| contrast | Log Odds 95% HDI | | | Log Odds 90% HDI | | | Second Factor 95% HDI | | Second Factor 90% HDI | |
| --- | --- | --- | --- | --- | --- | --- | --- | --- | --- | --- |
|  | Low | High | Sig. | Low | High | Sig. | Low | High | Low | High |
| Dwell: white  (ring − full) | 0.61 | 2.68 | * | 0.78 | 2.53 | * | 1.84 | 14.53 | 2.19 | 12.56 |
| Dwell: yellow  (ring − full) | 0.16 | 2.01 | * | 0.31 | 1.86 | * | 1.17 | 7.47 | 1.37 | 6.45 |
| Dwell: UV=null  (yellow − white) | -0.80 | 0.97 | x | -0.66 | 0.83 | x | 0.45 | 2.63 | 0.52 | 2.29 |
| Dwell: white  (ring − null) | -0.11 | 1.68 | x | 0.02 | 1.51 | * | 0.89 | 5.37 | 1.02 | 4.50 |
| Dwell: yellow  (ring − null) | 0.01 | 1.86 | * | 0.17 | 1.71 | * | 1.01 | 6.46 | 1.18 | 5.56 |

Table 2 Comparison between log odds of dwelling time within different fixed effect combination at 95% HDI and 90% HDI

The third time-spline (representing the smooth trend over the midday quartile) was significant at the 95% HDI in both the foraging and dwelling models, increasing the probability of foraging by ~87% and extending dwell times by approximately 3.1-fold. The second time-spline (spanning late morning into midday) reached significance only in the dwelling model (95% HDI), where it shortened dwell times to ~0.7-fold of baseline. The remaining spline terms crossed zero in both the 90% and 95% HDIs and were therefore considered non-significant. Notably, the monotone fixed effect of experimental day (the chronological replicate/learning effect) was also non-significant in both models.

| Time-spline | Log Odds 95% HDI | | | Log Odds 90% HDI | | | Odds Ratio 95% HDI | | Odds Ratio 90% HDI | |
| --- | --- | --- | --- | --- | --- | --- | --- | --- | --- | --- |
|  | Low | High | Sig. | Low | High | Sig. | Low | High | Low | High |
| 1 | -1.12 | 2.21 | x | -0.90 | 2.04 | x | 0.33 | 9.08 | 0.41 | 7.68 |
| 2 | -2.60 | 0.74 | x | -2.42 | 0.55 | x | 0.07 | 2.09 | 0.08 | 1.73 |
| 3 | 1.05 | 3.96 | * | 1.29 | 3.84 | * | 2.85 | 52.56 | 3.62 | 46.58 |
| 4 | -0.89 | 2.19 | x | -0.75 | 1.95 | x | 0.41 | 8.93 | 0.47 | 7.03 |

Table 3 Comparison between log odds of time-splines coefficient on foraging probability at 95% HDI and 90% HDI

| Time-spline | Log Odds 95% HDI | | | Log Odds 90% HDI | | | Second Factor 95% HDI | | Second Factor 90% HDI | |
| --- | --- | --- | --- | --- | --- | --- | --- | --- | --- | --- |
|  | Low | High | Sig. | Low | High | Sig. | Low | High | Low | High |
| 1 | -1.12 | 1.25 | x | -0.99 | 1.09 | x | 0.33 | 3.48 | 0.37 | 2.98 |
| 2 | -2.37 | -0.14 | * | -2.22 | -0.26 | * | 0.09 | 0.87 | 0.11 | 0.77 |
| 3 | 0.12 | 2.14 | * | 0.22 | 2.01 | * | 1.13 | 8.52 | 1.25 | 7.44 |
| 4 | -0.83 | 1.15 | x | -0.76 | 0.98 | x | 0.44 | 3.16 | 0.47 | 2.67 |

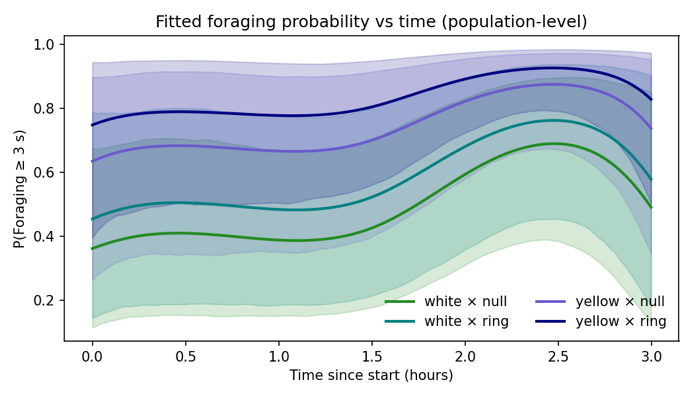

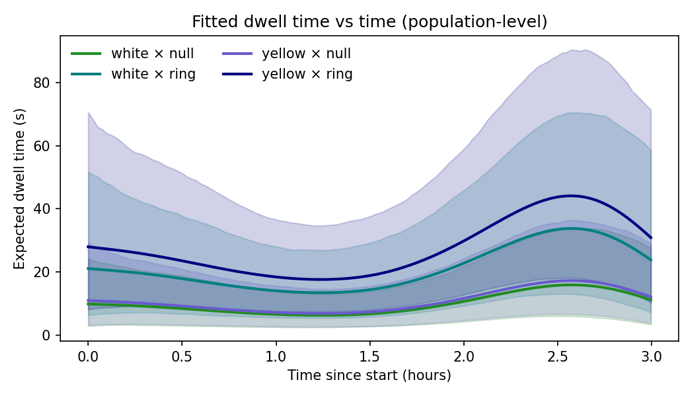

Table 4 Comparison between log odds time-splines coefficient on foragers dwelling time at 95% HDI and 90% HDI

From the z-standardized weather covariates, global radiation, downwelling radiation, and temperature at 2 m were all significant predictors in the foraging model (95% HDI). Global and downwelling radiation increased the odds of foraging by approximately 68% (OR ≈ 1.68) and 44% (OR ≈ 1.44), respectively, while higher temperature reduced the odds by about 34% (OR ≈ 0.66) relative to the model’s intercept. The same covariates showed consistent effects in the dwelling model (95% HDI): global and downwelling radiation prolonged dwelling time by about 1.27-fold and 1.53-fold, respectively, whereas temperature shortened it to about 0.63-fold of the baseline. Additionally, relative humidity showed marginal significance at the 90% HDI, reducing dwelling time to approximately 0.80-fold of the baseline.

Figure 5 Distribution of foraging probability (right) and dwelling time (left) among the four overlapping time-splines

| Parameter | Log Odds 95% HDI | | | Log Odds 90% HDI | | | Odds Ratio 95% HDI | | Odds Ratio 90% HDI | |
| --- | --- | --- | --- | --- | --- | --- | --- | --- | --- | --- |
|  | Low | High | Sig. | Low | High | Sig. | Low | High | Low | High |
| Downwelling radiation | 0.08 | 0.66 | * | 0.11 | 0.62 | * | 1.09 | 1.94 | 1.12 | 1.85 |
| Global radiation | 0.10 | 0.90 | * | 0.17 | 0.87 | * | 1.10 | 2.47 | 1.18 | 2.40 |
| Relative humidity | -0.85 | 0.12 | x | -0.76 | 0.08 | x | 0.43 | 1.12 | 0.47 | 1.09 |
| Temperature | -1.80 | -0.40 | * | -1.71 | -0.48 | * | 0.17 | 0.67 | 0.18 | 0.62 |
| Wind speed | -0.45 | 0.06 | x | -0.43 | 0.01 | x | 0.64 | 1.06 | 0.65 | 1.01 |

Table 5 Comparison between log odds of weather coefficients on foraging probability at 95% HDI and 90% HDI

| Parameter | Log Odds 95% HDI | | | Log Odds 90% HDI | | | Second Factor 95% HDI | | Second Factor 90% HDI | |
| --- | --- | --- | --- | --- | --- | --- | --- | --- | --- | --- |
|  | Low | High | Sig. | Low | High | Sig. | Low | High | Low | High |
| Downwelling radiation | 0.27 | 0.59 | * | 0.29 | 0.57 | * | 1.31 | 1.80 | 1.33 | 1.76 |
| Global radiation | 0.02 | 0.47 | * | 0.04 | 0.43 | * | 1.02 | 1.60 | 1.04 | 1.54 |
| Relative humidity | -0.49 | 0.02 | x | -0.44 | -0.001 | * | 0.61 | 1.01 | 0.64 | 1.00 |
| Temperature | -0.89 | -0.07 | * | -0.84 | -0.1 | * | 0.41 | 0.93 | 0.43 | 0.89 |
| Wind speed | -0.15 | 0.10 | x | -0.14 | 0.09 | x | 0.86 | 1.10 | 0.87 | 1.09 |

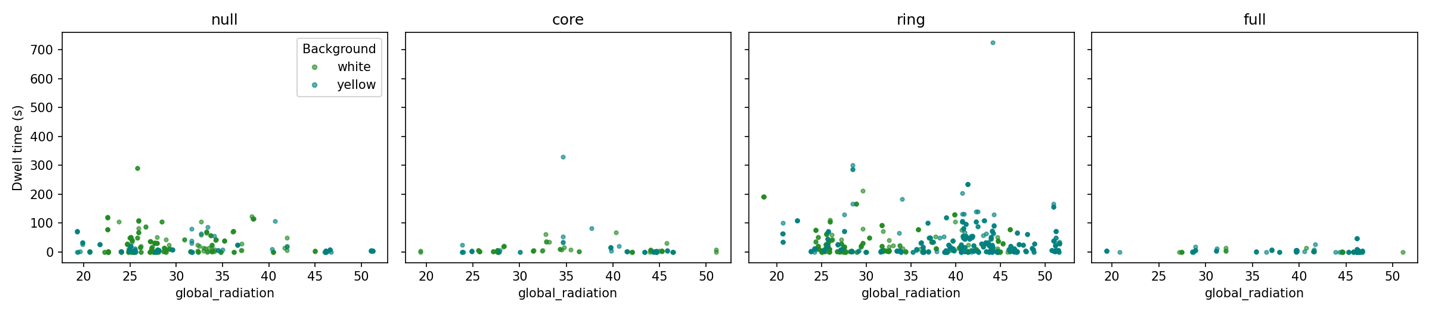
Table 6 Comparison between log odds of weather coefficients on foragers dwelling time at 95% HDI and 90% HDI

Figure 6 Effect of global radiation values within different UV patch allocations and artificial floral ground colour on foragers dwelling time

Finally, variance partitioning of the random effects revealed that in the foraging model, ~52% of the variance was attributable to FlowerID, ~2% to observer bias, and only ~0.65% to bee identity, indicating strong homogeneity among individuals. The remaining ~43% of variance was captured by the latent residual component. A similar pattern was observed for the dwelling model, with ~31% of the variance explained by FlowerID, ~2% by observer bias, and ~0.75% by bee identity, while more than 64% remained in the residual. This substantial residual share suggests an underlying complexity and stochasticity in pollinator behaviour beyond the measured effects.

| Model | component | median | 95% HDI | | 90% HDI | |
| --- | --- | --- | --- | --- | --- | --- |
|  |  |  | Low | High | Low | High |
| Foraging (logit) | bee | 0.651077 | 0.001854 | 3.960744 | 0.006832 | 3.091789 |
| Foraging (logit) | flower | 52.651524 | 37.732571 | 66.464351 | 40.109523 | 64.280404 |
| Foraging (logit) | observer | 2.115150 | 0.150194 | 16.448600 | 0.279568 | 12.368981 |
| Foraging (logit) | residual(latent) | 42.857113 | 30.049397 | 56.256267 | 32.197699 | 54.086900 |
| Dwell (log-normal AFT) | bee | 0.750330 | 0.006093 | 2.790531 | 0.021280 | 2.415506 |
| Dwell (log-normal AFT) | flower | 31.333927 | 21.553743 | 41.980139 | 23.182908 | 40.274669 |
| Dwell (log-normal AFT) | observer | 1.943382 | 0.177857 | 20.941142 | 0.296670 | 15.129894 |
| Dwell (log-normal AFT) | residual(log-scale) | 64.162447 | 50.080377 | 74.021038 | 53.189681 | 72.650947 |

Table 7 Standard deviation of noise for random effects and models reiduals at )5% HDI and 90% HDI
